# WormCat: an online tool for annotation and visualization of *Caenorhabditis elegans* genome-scale data

**DOI:** 10.1101/844928

**Authors:** Amy D. Holdorf, Daniel P. Higgins, Anne C. Hart, Peter R. Boag, Gregory J. Pazour, Albertha J. M. Walhout, Amy K. Walker

**Affiliations:** Program in Systems Biology, University of Massachusetts Medical School, Worcester, MA 01605, USA; Department of Computer Science, Georgia Technical University, Atlanta, GA 30332-0765, USA; Department of Neuroscience, Robert J. and Nancy D. Carney Institute for Brain Science, Brown University, Providence, RI 02912, USA; Department of Biochemistry and Molecular Biology, Monash University, 3800 Clayton Australia; Program in Molecular Medicine, University of Massachusetts Medical School, Worcester, MA 01605, USA

## Abstract

The emergence of large gene expression datasets has revealed the need for improved tools to identify enriched gene categories and visualize enrichment patterns. While Gene Ontogeny (GO) provides a valuable tool for gene set enrichment analysis, it has several limitations. First, it is difficult to graphically compare multiple GO analyses. Second, genes from some model systems are not well represented. For example, around 30% of *Caenorhabditis elegans* genes are missing from analysis in commonly used databases. To allow categorization and visualization of enriched *C. elegans* gene sets in different types of genome-scale data, we developed WormCat, a web-based tool that uses a near-complete annotation of the *C. elegans* genome to identify co-expressed gene sets and scaled heat map for enrichment visualization. We tested the performance of WormCat using a variety of published transcriptomic datasets and show that it reproduces major categories identified by GO. Importantly, we also found previously unidentified categories that are informative for interpreting phenotypes or predicting biological function. For example, we analyzed published RNA-seq data from *C. elegans* treated with combinations of lifespan-extending drugs where one combination paradoxically shortened lifespan. Using WormCat, we identified sterol metabolism as a category that was not enriched in the single or double combinations but emerged in a triple combination along with the lifespan shortening. Thus, WormCat identified a gene set with potential phenotypic relevance that was not uncovered with previous GO analysis. In conclusion, WormCat provides a powerful tool for the analysis and visualization of gene set enrichment in different types of *C. elegans* datasets.

## Introduction

RNA-seq is an indispensable tool for understanding how gene expression changes during development or upon environmental perturbations. As this technology has become less expensive and more robust, it has become more common to generate data from multiple conditions, enabling comparisons of gene expression profiles across biological contexts. The most commonly used method to derive information on the biological function of co-expressed genes is Gene Ontology (GO) (The Gene Ontology 2019) (Ashburner *et al*. 2000), where each gene has been annotated by three major classifications: *Biological Process*, *Molecular Function* or *Cellular Component*. For example, the *Biological Process* class is defined as a process that an organism is programmed to execute, and that occurs through specific regulated molecular events. *Molecular Function* refers to protein activities, and *Cellular Component* maps the location of activity. Within each of these classifications, functions are broken down in parent-child relationships with increasing functional specificity (**Fig 1A**). However, child classes can be linked to different parent classes, making statistical analysis not straightforward. For example, the child class *phospholipid biosynthetic process* can be linked to both of the parent groups *metabolic process* and *cellular process*. Thus, GO provides multiple descriptors per gene. Although GO was developed to compare gene function across newly sequenced genomes, it became apparent that it could also be used to identify shared functional classifications within large-scale gene expression data (Eisen *et al*. 1998; Spellman *et al*. 1998). Currently, multiple web-based servers that use different statistical tests can be used to determine enrichment of GO terms for a gene set of interest. For example, PANTHER (www.pantherdb.org) provides enriched GO terms determined by Fisher’s Exact test with a Benjamini-Hochberg false discovery rate (FDR) correction for 131 species (Mi *et al*. 2019). Because the multiplicity of GO term parent-child relationships can produce complex data structures, specialized ontologies such as GO-Slim use a restricted set of terms, searching biological processes as default (Mi *et al*. 2019). *P-*values are provided for enriched GO terms. Visualization of gene set enrichment data is important for identifying critical elements and communication of information. PANTHER provides pie or bar charts of individual searches (Mi *et al*. 2019). The GOrilla platform generates tables of *P-*values (Eden *et al*. 2009) and links to another service, REVIGO, that use semantic graphs to visualize GO terms data (Supek *et al*. 2011). Thus, the GO databases provide a widely used platform for classifying, comparing, and visualizing functional genomic data. However, as outlined below, GO is of limited use for the analysis of *Caenorhabditis elegans* data and visualization of multiplexed datasets.

**Figure 1:**
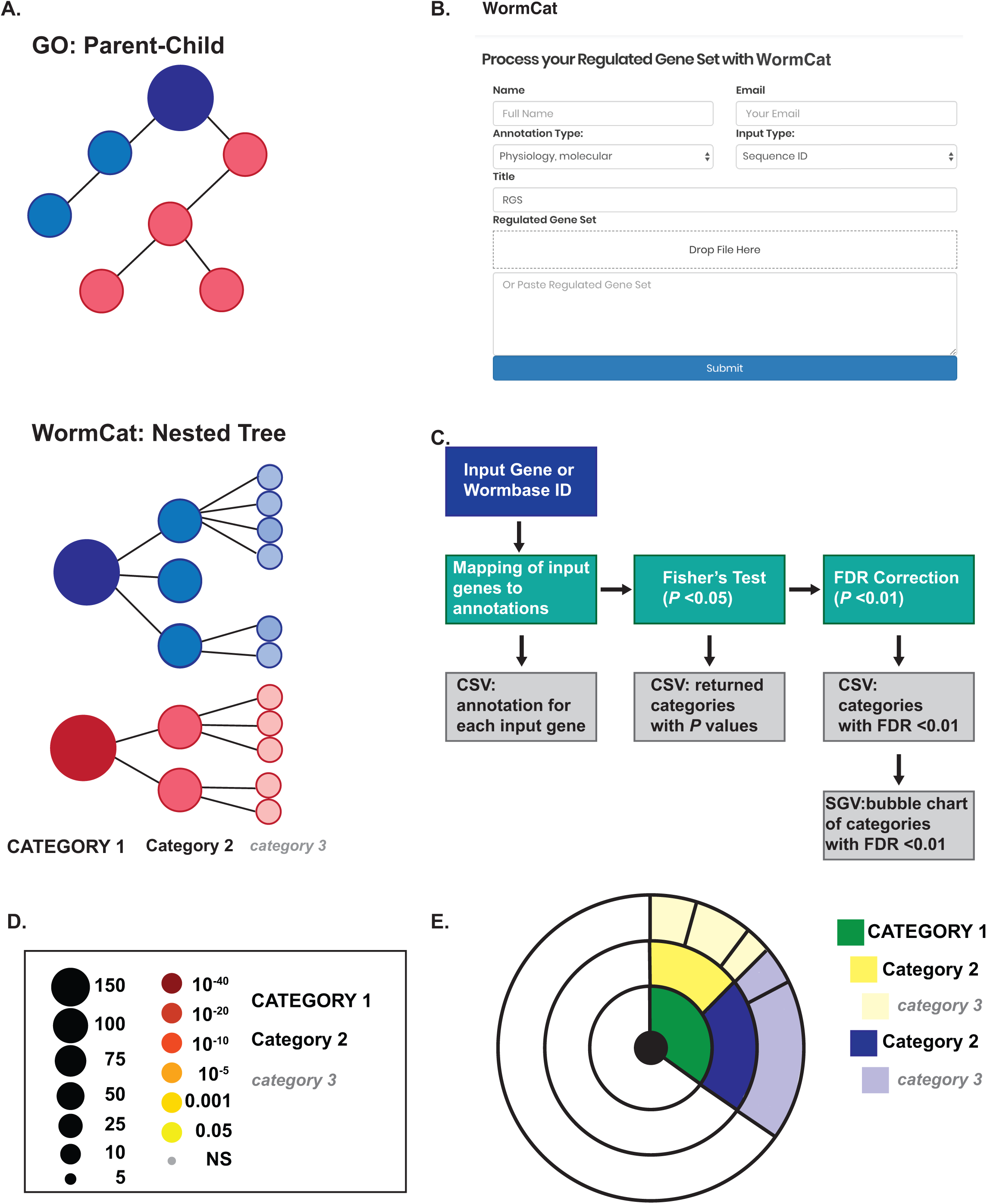
WormCat annotates and visualizes *C. elegans* gene enrichment from genome-scale data. (**A**) Diagram comparing the parent-child methods for linking GO terms with the nested tree strategy used for annotating *C. elegans* genes in WormCat. (**B**) Screenshot of the WormCat web page showing the data entry form. (**C**) Flow chart diagraming steps and outputs from the WormCat program. Data outputs are in tabular comma-separated values (CSV) and scalable vector graphics (SVG) formats. (**D**) Legend for scaled bubble charts showing number of genes referenced to size and *P-*value referenced to color. In graphs, Category 1, 2 and 3 are differentiated by capitalization, size and italics. (**E**) Legend for sunburst plots showing concentric rings visualizing Category 1, 2 and 3 data.

The nematode *C. elegans* has been at the forefront of genomics research. It was the first metazoan organism with a completely sequenced genome (Consortium 1998). After the discovery of RNA interference (RNAi)(Fire *et al*. 1998), multiple RNAi libraries were developed for performing genome-wide knockdown screens (Kamath *et al*. 2003; Rual *et al*. 2004). Gene expression profiling studies using microarrays or RNA-seq have compared gene expression in sex-specific, developmental/aging-related, specific gene deletion, tissue-specific, and dietary or stress-related animal conditions (Reinke *et al*. 2000; Hillier *et al*. 2005; Baugh *et al*. 2009; Oliveira *et al*. 2009; Deng *et al*. 2011; Schwarz *et al*. 2012; Bulcha *et al*. 2019). While GO has been used extensively to analyze *C. elegans* gene expression profiling data, it has several limitations. First, around 30% of *C. elegans* genes are not annotated in GO databases (Ding *et al*. 2018), excluding these genes from analysis. Thus, these genes are arbitrarily excluded from enrichment statistics. Second, the visualization of enrichment data from comparative RNA-seq datasets is difficult, and this is true not only for *C. elegans* datasets, but for gene expression profile comparisons in any organism. Most users display the output data as lists with *P-*values (Macneil *et al*. 2013) or as pie or bar charts (Ding *et al*. 2015), which are not easily multiplexed for comparison of multiple datasets. Finally, it can be challenging to determine which input genes are associated with a given GO classification, which is critical for interpreting the accuracy and biological importance of enriched gene sets.

We constructed a web-based gene set enrichment analysis tool we named WormCat (WormCatalog) that works independently from GO to identify potentially co-expressed or co-functioning genes in genome-wide expression studies or functional screens. WormCat (www.wormcat.com), uses a concise list of nested categories where each gene is first assigned to a category based on physiological function, and then to a molecular function or cellular location. WormCat provides a scaled bubble chart that allows the visualization and direct comparison of complex datasets. The tool also provides csv files containing input gene annotations, *P-*values from Fisher’s exact tests and Bonferroni multiple hypothesis testing corrections. We used WormCat to identify functional gene sets in published gene expression data and large-scale RNAi screens. WormCat reproducibly identified prior GO classifications and provided an easy way to interpret visualization that enables the facile and intuitive comparison of multiple published datasets. We also identified new groups of enriched categories with potentially important biological significance, showing that WormCat provides enrichment information not revealed by GO. Taken together, WormCat offers an alternative and complementary tool for categorizing and visualizing data for genome-wide *C. elegans* studies and may provide a platform for similar annotations in other model organisms and humans.

## Materials and Methods

### Annotations

WormBase version WS270 was used to provide WormBase descriptions and provide phenotype information.

### Scripts

The processed data were analyzed using R version 3.4.4 (2018-03-15) and depends on the following R packages: datasets, graphics, grDevices, methods, stats, utils, ggplot2, plot flow, scales, ggthemes, pander, data.table, plyr, gdtools, svglite, FSA.

### Data Availability

The code and annotation lists are available under MIT Open Source License and can be downloaded from the GitHub repository https://github.com/dphiggs01/wormcat along with version-control information. Alternatively, WormCat can be installed directly as an R package using the devtools library. Supplemental material has been deposited at fig share and includes twelve supplemental figures and fourteen supplemental tables.

### GO searches

Genes lists were entered as test sets into GOrilla (http://cbl-gorilla.cs.technion.ac.il/) (Eden *et al*. 2009) with the WormCat annotation list used as background so that the same background set was used when comparing WormCat and GOrilla. *All* was selected for ontogeny choices and the *P*-value thresholds were set to 10^−3^. Output selections were Microsoft Excel and REVIGO (Supek *et al*. 2011).

## Summary of Supplementary Figures

**Figure S1. GO Analysis of upregulated genes from *sams-1(RNAi)* animals by Gene Set Enrichment Analysis.** Bar graph showing GO categories returned from *sams-1(RNAi)* upregulated genes (Ding *et al*. 2015) by the WormBase Gene Set Enrichment Analysis tool (Angeles-Albores *et al*. 2016).

**Figure S2. WormCat verifies known category enrichments from *sams-1(RNAi)* downregulated genes**. (**A-B**) Semantic graphs of GO analysis generated by GOrilla (Eden *et al*. 2009) and visualized by REVIGO (Supek *et al*. 2011) of *sams-1(RNAi)* downregulated genes from untreated (**A**) and choline (Ch) treated (**B**) animals. (**C**) WormCat bubble heat plots comparing *sams-1(RNAi)* with and without choline. Gene expression microarray data for **A-C** were obtained from Ding *et al*., 2015. Bubble heat plot key is the same as **Fig 1D**. CUB, Complement C1r/C1s, Uegf, Bmp1 Domain; PUF, Pumilio and *fem-3* mRNA Binding Factor; ZF, Zinc Finger. (**D**) Venn diagrams showing overlap between *Stress Response* genes in *sams-1(RNAi)* up (pink) or downregulated genes (blue).

**Figure S3: WormCat analysis of germline-specific microarray data identifies the tau tubulin kinase family as a male-specific category**. (**A**) Category 1 analysis of Oogenic (Oo) or Spermatogenic (Sp) data sets ordered by most enriched in Oo data. Breakdown of data from the Category 1 level for Cell cycle (**B**), Development (**C**), mRNA Functions (**D**), or Cytoskeleton (**E**). All data is from Reinke *et al*. Gen. Trans. Machinery, General Transcription Machinery; Trans. Chromatin, Transcription: Chromatin; ZF, zinc finger.

**Figure S4: GO analysis visualized by REVIGO of germline RNA-seq data from Ortiz *et al*.** Semantic graphs of GO analysis generated by GOrilla(Eden *et al*. 2009)and visualized by REVIGO of Gender Neutral (**A**), Oogenic (**B**) and Spermatogenic (**C**) germlines.

**Figure S5: GO analysis visualized by REVIGO of germline microarray data from Reinke *et al*.** Semantic graphs of GO analysis generated by GOrilla and visualized by REVIGO of Oogenic (**A**) and Spermatogenic (**B**) germlines.

**Figure S6: GO analysis visualized by REVIGO of larval tissue specific microarray data from Spencer *et al*.** Semantic graphs of GO analysis generated by GOrilla and visualized by REVIGO of microarray from larval Muscle (BWM, body wall muscle) (**A**) Intestine (Int) (**B**), Hypodermis (Hyp) (**C**), Excretory cells (Exc) (**D**), or Neurons (**E**).

**Figure S7: GO analysis visualized by REVIGO of adult tissue specific RNA-seq data from Kaletsky, *et al*.** Semantic graphs of GO analysis generated by GOrilla and visualized by REVIGO of RNA-seq data from adult Muscle (Mus) (**A**) Intestine (Int) (**B**), Hypodermis (Hyp) (**C**), or Neurons (**D**).

**Figure S8: GO analysis visualized by REVIGO of larval neuronal subtype microarray data from Spencer, et al.** Semantic graphs of GO analysis generated by GOrilla and visualized by REVIGO of microarray from larval dopaminergic (Dopa) (**A**) GABAergic (GABA) (**B**), *glr-1* expressing (*glr-1*) (**C**), or Class A motor neurons (Motor) (**D**).

**Figure S9: WormCat analysis of upregulated genes in *C. elegans* treated with triple combinations of lifespan-changing drugs**. Category 1, 2, and 3 analysis of upregulated genes found by RNA-seq from triple-drug combinations (Admasu *et al*. 2018). Pink box denotes drug combination that causes premature death. Allan, allantoin; CYP, Cytochrome P450; EC Material, Extracellular Material; Maj Sperm Protein, Major Sperm Protein; Neur Function; Neuronal Function; NHR, Nuclear Hormone Receptor; Prot General, Proteolysis General; Psora, Psora-4; Rapa, Rapamycin; Rifa, Rifampicin; Short Chain Dehydr., Short Chain Dehydrogenase; Trans Factor, Transcription Factor; TYR kinase, Tyrosine Kinase; ugt, UDP-glycosyltransferase.

**Figure S10: WormCat analysis of downregulated genes in *C. elegans* treated with triple combinations of lifespan-changing drugs**. Category 1, 2, and 3 analysis of downregulated genes found by RNA-seq from triple-drug combinations (Admasu *et al*. 2018). Pink box denotes drug combination that causes premature death. Allan, Allantoin; Psora, Psora-4; Rapa, Rapamycin; Rifa, Rifampicin.

**Figure S11: GO analysis visualized by REVIGO of upregulated genes from RNA-seq data from *C. elegans* treated with triple combinations of lifespan extending drugs from Admasu *et al*.** Semantic graphs of GO analysis generated by GOrilla and visualized by REVIGO of RNA-seq data from Rifa, Psora, Allan treated (**A**) Rifa, Rapa, Allan treated (**B**), or Rifa, Rapa, Psora treated (**C**).

**Figure S12: GO analysis visualized by REVIGO of RNA-seq data from a *C. elegans* RNAi screen for glycogen storage from LaMacchia *et al*.** Semantic graphs of GO analysis generated by GOrilla and visualized by REVIGO of *C. elegans* showing low glycogen storage in an RNAi screen.

## Summary of Supplementary Tables

**Supplemental Table 1: WormCat annotations.** xlsx file containing *C. elegans* genes arranged alphabetically by Categories with Sequence ID, WormBase ID, and Category 1, 2, and 3 annotations along with WormBase descriptions (WormBase version WS270).

**Supplemental Table 2: WormCat annotation definitions**. xlsx file containing annotation definitions.

**Supplemental Table 3: Random gene analysis.** xlsx file with tabs containing lists of randomly generated WormBase IDs of 100 (tabs 1-4), 500 (tabs 5-8), 1000 (tabs 9-12 and 1500 (tabs 13-16) genes with Category 1, 2 and 3 analysis. NA genes on these tables reflect WormBase IDs that have been merged, marked as dead or updated as not corresponding to a protein-coding gene.

**Supplemental Table 4: GO analysis of *sams-1(RNAi)* regulated genes.** xlsx file containing GO terms produced by GOrilla (Eden *et al*. 2009) from microarray data for *sams-1(RNAi)* up genes (tab 1), *sams-1(RNAi)* up plus choline (CH) (tab 2), *sams-1(RNAi)* down (tab 3), *sams-1(RNAi)* down plus CH (tab 4), and the *sams-1(RNAi)* up genes identified by GOrilla as lipid metabolism with corresponding WormCat annotations (tab 5). Data from Ding *et al.*, 2015.

**Supplemental Table 5: WormCat analysis of *sams-1(RNAi)* regulated genes.** xlsx file containing Category 1, 2, and 3 analysis from microarray data for *sams-1(RNAi)* up and down genes with or without choline (CH, tabs 1-3) from Ding *et al*., 2015. Tabs 4-7 contain input genes with WormCat annotations for each gene. NA genes on these tables reflect WormBase IDs that have been merged, marked as dead or updated as not corresponding to a protein-coding gene.

**Supplemental Table 6: WormCat analysis of *sams-1(RNAi)* regulated genes that were excluded by GSEA analysis.** xlxs file containing Category 1, 2, and 3 analysis from microarray data for *sams-1(RNAi)* upregulated genes (see **Table S5**, Tab 4) that were excluded by the GSEA tool on WormBase. Data from Ding *et al*., 2015.

**Supplemental Table 7: WormCat analysis of germline-expressed genes from Oritz *et al*.** xlxs file containing Category 1, 2, and 3 analysis from RNA-seq data for Germline Neutral (GN), Oogenic (Oo) or Spermatogenic (Sp) datasets (tabs 1-3). Tabs 4-6 contain input genes with WormCat annotations for each gene. NA genes on these tables reflect WormBase IDs that have been merged, marked as dead or updated as not corresponding to a protein-coding gene. Tabs 7-10 contain GO analysis by GOrilla for Germline Neutral (GN), Oogenic (Oo) or Spermatogenic (Sp) datasets.

**Supplemental Table 8: WormCat analysis of germline-expressed genes from Reinke *et al*.** xlxs file containing Category 1, 2, and 3 analysis from microarray data of Oogenic (Oo) or Spermatogenic (Sp) datasets (tabs1-3). Tabs 4-5 contain input genes with WormCat annotations for each gene. NA genes on these tables reflect WormBase IDs that have been merged, marked as dead or updated as not corresponding to a protein-coding gene. Tabs 6-7 contain GO analysis by GOrilla for Oogenic (Oo) or Spermatogenic (Sp) datasets.

**Supplemental Table 9: WormCat analysis of larval tissue-specific genes from Spencer *et al*.** xlsx file containing Category 1, 2, and 3 analysis from microarray data from *selective enriched* datasets (tabs 1-3). Tabs 4-12 contain input genes with WormCat annotations for each gene for all tissue and cell types examined. NA genes on these tables reflect WormBase IDs that have been merged, marked as dead or updated as not corresponding to a protein-coding gene. Tabs 13-22 contain GO analysis by GOrilla from selective enriched datasets from Muscle (BWM, body wall muscle), Intestine (Int), Hypodermis (Hyp), Excretory cells (Exc), Neurons (Neuro, pan-neuronal), Dopaminergic (Dopa), GABAergic (GABA), *glr-1* expressing (*glr-1*) or Class A Motor neurons (Motor).

**Supplemental Table 10: WormCat analysis of adult tissue-specific genes from Kaletsky *et al*.** xlsx file containing Category 1, 2, and 3 analysis from RNA-seq data of enriched (en) and unique (un) datasets (tabs 1-3). Tabs 4-11 contain input genes with WormCat annotations for each gene for all tissue and cell types examined. NA genes on these tables reflect WormBase IDs that have been merged, marked as dead or updated as not corresponding to a protein-coding gene. Tabs 12-16 contain GO analysis by GOrilla of enriched genes from Muscle (Mus), Intestine (Int), Hypodermis (Hyp) or Neuron (Neur).

**Supplemental Table 11: WormCat analysis of upregulated genes from a combinatorial RNA-seq study of lifespan enhancing drugs.** xlsx file analysis of data from Admasu *et al*. comparing upregulated genes from single, double, and triple combinations of lifespan inducing drugs. Tabs 1-3: Category 1-3 analysis of genes upregulated by single drugs. Tabs 4-7: Input genes with WormCat annotations for each single drug treatment. Tabs 8-10: Category 1-3 analysis of upregulated genes by double drug combinations. Tabs 11-14: Input genes with WormCat annotations for each double combination drug treatment. Tabs 15-17: Category 1-3 analysis of upregulated genes by triple-drug combinations. Tab 18: Genes from the *Metabolism: lipid: sterol* category from the Rifa/Rapa/Psora set with corresponding lifespan data from Murphy *et al*. (yellow) (Murphy *et al*. 2003). Tabs 19-21: Input genes with WormCat annotations for each triple combination drug treatment. NA genes on these tabs reflect WormBase IDs that have been merged, marked as dead or updated as not corresponding to a protein-coding gene. Tabs 22-24: GO analysis by GOrilla (Eden *et al*. 2009) for each triple combination drug treatment.

**Supplemental Table 12: WormCat analysis of downregulated genes from a combinatorial RNA-seq study of lifespan enhancing drugs.** xlsx file analysis of data from Admasu *et al*. comparing downregulated genes from single, double and triple combinations of lifespan inducing drugs. Tabs 1-3: Category 1-3 analysis of genes downregulated by single drugs. Tabs 4-7: Input genes with WormCat annotations for each single drug treatment. Tabs 8-10: Category 1-3 analysis of downregulated genes by double drugs combinations. Tabs 11-14: Input genes with WormCat annotations for each double combination drug treatment. Tabs 15-17: Category 1-3 analysis of downregulated genes by triple-drug combinations. Tabs 18-20: input genes with WormCat annotations for each triple combination drug treatment. NA genes on these tabs reflect WormBase IDs that have been merged, marked as dead or updated as not corresponding to a protein-coding gene.

**Supplemental Table 13: WormCat annotations for genes in the Ahringer RNAi library**. xlsx file containing gene and WormBase IDs along with Category 1, 2, and 3 annotations for the clones represented in the Ahringer RNAi library (Kamath *et al*. 2003).

**Supplemental Table 14: WormCat analysis of a genome scale RNAi screen from LaMacchia *et al*.** xlsx file containing Category 1, 2, and 3 analysis of RNAi screen data of all, high and low glycogen stained animals (Tabs 1-3). Tabs 4-6 contain input genes with WormCat annotations for each gene. Tab 7 is Glycogen low genes analyzed by GOrilla. Tabs 8-11 show GO terms with multiple categories containing *cyc-1*, *vha-6*, *pbs-7* and Y71F9AL.17.

## Results

### *C. elegans* gene annotation

The *C. elegans* genome encodes ~19,800 protein-coding genes, ~260 microRNAs and numerous other non-coding RNAs (WormBase version WS270). We annotated all *C. elegans* genes first based on physiological functions, and, when these functions were unknown or pleiotropic, according to molecular function or sub-cellular location (See **Table S1** for annotations, **Table S2** for Category definitions). Our annotations are structured as nested categories, enabling classification into broad (Category 1; Cat1), or more specific categories (Category 2 or 3; Cat2 or Cat3). This annotation has the advantage of including information from multiple sources in addition to GO. For example, we used phenotype information available in WormBase (Lee *et al*. 2018), for Cat1 assignments. Importantly, the phenotypic data present in WormBase (Lee *et al*. 2018) was only used if phenotypes were: 1) derived from wild type animals, 2) examined in detail in peer-reviewed publications, and (?) 3) represented in two independent screens. If a gene was ascribed a clear physiological function with these criteria, we assigned it to a physiological category, examples of which include *Stress response*, *Development*, and *Neuronal function*. If gene products have multiple functions within the cell, act in multiple cells type or different developmental times, we prioritized assignment to molecular categories. Molecular categories harbor both genes whose products comprise molecular machines, as well as the chaperones or regulatory factors that are necessary for the function of such machines. We used information on molecular function of human orthologs to classify *C. elegans* genes that had not been molecularly defined in nematodes and showed highly similar in BLAST scores. For example, we classified the *C. elegans* gene W03D8.8 in *Metabolism: lipid: beta oxidation* based on a BLAST score of *e* = 7×10^−37^ and similarity over 92% of its length to human ACOT4 (acyl-CoA thioesterase 4). For genes with weaker homology to human genes, we further refined assignments using BLAST (Altschul *et al*. 1990) and the NCBI Conserved Domain server (Marchler-Bauer *et al*. 2017). We used these tools to determine if there was significant homology or shared domains between *C. elegans* and human proteins, then used information in UniProt (www.uniprot.org) for the human proteins to determine molecular classification. For example, we placed the *C. elegans* gene T26E4.3 in *Protein modification: carbohydrate* based on a BLAST core of *e* = 4×10^−7^ over 95% of its length to human alpha fucosyltransferase 1 and identification of a Fut1_Fut2-like domain by the NCBI conserved domain server with an *e* score of 6.16×10^−36^. However, while the gene BE10.3 is referred to in the WormBase description as an ortholog of human FUT9 (fucosyltransferase 9) (**Table S1**), we found no homology to human genes by NCBI BLAST or domain conservation across all organisms with the NCBI Conserved Domain server. Therefore, we classified BE10.3 in *Unknown*. Finally, if no biological or molecular function could be assigned, protein sub-cellular localization was used for annotation. For example, a protein with a predicted membrane-spanning region that lacks characterization as a receptor would be placed in *Transmembrane protein*. Genes with no functional information were classified as *Unknown* (Cat1). There are 8160 genes that lacked sufficient information for classification in physiological, molecular or sub-cellular localization categories and were classified in *Unknown*. Many of these genes are *C. elegans* or nematode-specific, however, some have homology to human genes of unknown function. WormBase also aggregates microarray and RNA-seq information and annotates genes that respond to pharmacological treatments (Lee *et al*. 2018). We also used this information to differentiate genes within *Unknown: regulated by multiple stresses* that respond to at least two commonly used stressors. This classification does not imply these genes have a function in the stress response. It does allow identification of genes with otherwise unknown functions that are common responders to stress. This may be useful to distinguish RNA-seq datasets that respond similarly to pharmacological stressors or can serve as a source to identify specific genes of interest for additional study. We also included pseudogenes and non-coding RNAs in our annotation list. These genes commonly appear in RNA-seq data; including them in the annotation list allows them to be labeled within the user’s input dataset. In this way, we were able to leverage multiple data sources to categorize *C. elegans* genes into potentially functional biological groups.

### WormCat.com allows web-based searches of input genes and generates scaled bubble charts and gene lists

WormCat.com maps annotations to input genes, then determines category enrichment for Cat1, Cat2 and Cat3 (**Fig 1B**). Determination of category enrichment in a gene set of interest compared to the entire genome can rely on several commonly used statistics such as the Fisher’s exact test and the Mann-Whitney test (Mi *et al*. 2019). We used Fisher’s exact test to determine if categories were overrepresented because it is accurate down to small sample sizes, which may occur in high resolution classifications (Mcdonald 2014). In addition, we included the Bonferroni FDR correction (Mcdonald 2014). To determine the number of false positives after the Fisher’s test or the FDR correction, we tested randomized gene lists of 100, 500, 1000, 1500 genes and found that small numbers of genes were returned using a *P-*value cut-off of 0.05 (for, example 5 genes were returned on the 1000 gene random set). Few genes were returned from any of the randomized sets using an FDR cutoff of 0.01 (**Table S3**). Because an FDR < 0.01 is relatively stringent, Fisher’s exact test *P*-values will also be provided allowing users to make independent evaluations on the statistical cut-offs.

The WormCat website (www.wormcat.com) provides gene enrichment outputs in multiple formats (**Fig 1C**). First, all input genes are listed with mapped annotations (rgs_and_categories.csv). Genes that matched at least one Cat1, Cat2 and Cat3 classification are returned with Fisher’s exact test *P-*values (Cat1.csv, Cat2.csv or Cat3.csv). Next, Cat1, Cat2, and Cat3 matches with an FDR correction of < 0.01 are returned as CSV files named Cat1.apv, Cat2.apv and Cat3.apv (appropriate *P-*value). Finally, the Cat.apv files are used to generate two types of graphical output. First, it constructs scaled heat map bubble charts (Cat1., Cat2., Cat3.sgv) where color signifies *P-*values and size specifies the number of genes in the category (**Fig 1D**). The scaling for these graphs is fixed so that multiple datasets can be compared and graphed together. Second, a sunburst graph is built with concentric rings of Cat1, Cat2, and Cat3 values (**Fig 1E**). In these graphs, sections of each ring correspond to categories, with the size of the section proportional to the number of genes in the category. On the website, each ring section is clickable to generate a sub-graph-based division within a section. For example, clicking a single Cat1 section would generate a subgraph with all the Cat2 and Cat3 subdivisions located within. This graphical output is likely to be most useful for visualization of a single RNA-seq dataset, or genetic screening data. Thus, WormCat provides multiple outputs to allow inspection of individual input genes, generation of gene tables and *P-*values, and graphical visualization of enrichments.

### Comparison of GO and WormCat analysis of *sams-1(RNAi)* enrichment data

To determine the utility of the WormCat annotations, we first analyzed microarray data we previously generated to compare gene expression changes after knockdown of *sams-1*, with and without dietary supplementation of choline (Ding *et al*. 2015). *sams-1* encodes an S-adenosylmethionine (SAM) synthase, which is an enzyme that produces nearly all of the methyl groups used in methylation of histones and nucleic acids, in addition to the production of the membrane phospholipid phosphatidylcholine (PC) (Mato and Lu 2007). *sams-1* RNAi or loss-of-function *(lof)* animals have extended lifespan (Hansen 2005), increased lipid stores (Walker *et al*. 2011), and activated innate immune signatures (Ding *et al*. 2015). *sams-1* animals have low PC (Walker *et al*. 2011), but those levels are restored with supplementation of choline (Ding *et al*. 2015), which supports SAM-independent phosphatidylcholine synthesis (Vance 2014) (**Fig 2A**). Gene expression changes in *sams-1(RNAi)* animals could result from perturbation in different SAM-dependent pathways. To determine which transcriptional changes occurred downstream of alterations in PC synthesis, we performed microarrays with RNA from *sams-1(RNAi)* and *sams-1(RNAi)* animals supplemented with choline. 90% of genes that changed in expression in *sams-1(RNAi)* animals returned to wild type levels after choline supplementation. Therefore, the expression of the remaining 10% of genes was altered by *sams-1* RNAi independently of phosphatidylcholine levels (Ding *et al*. 2015).

**Figure 2:**
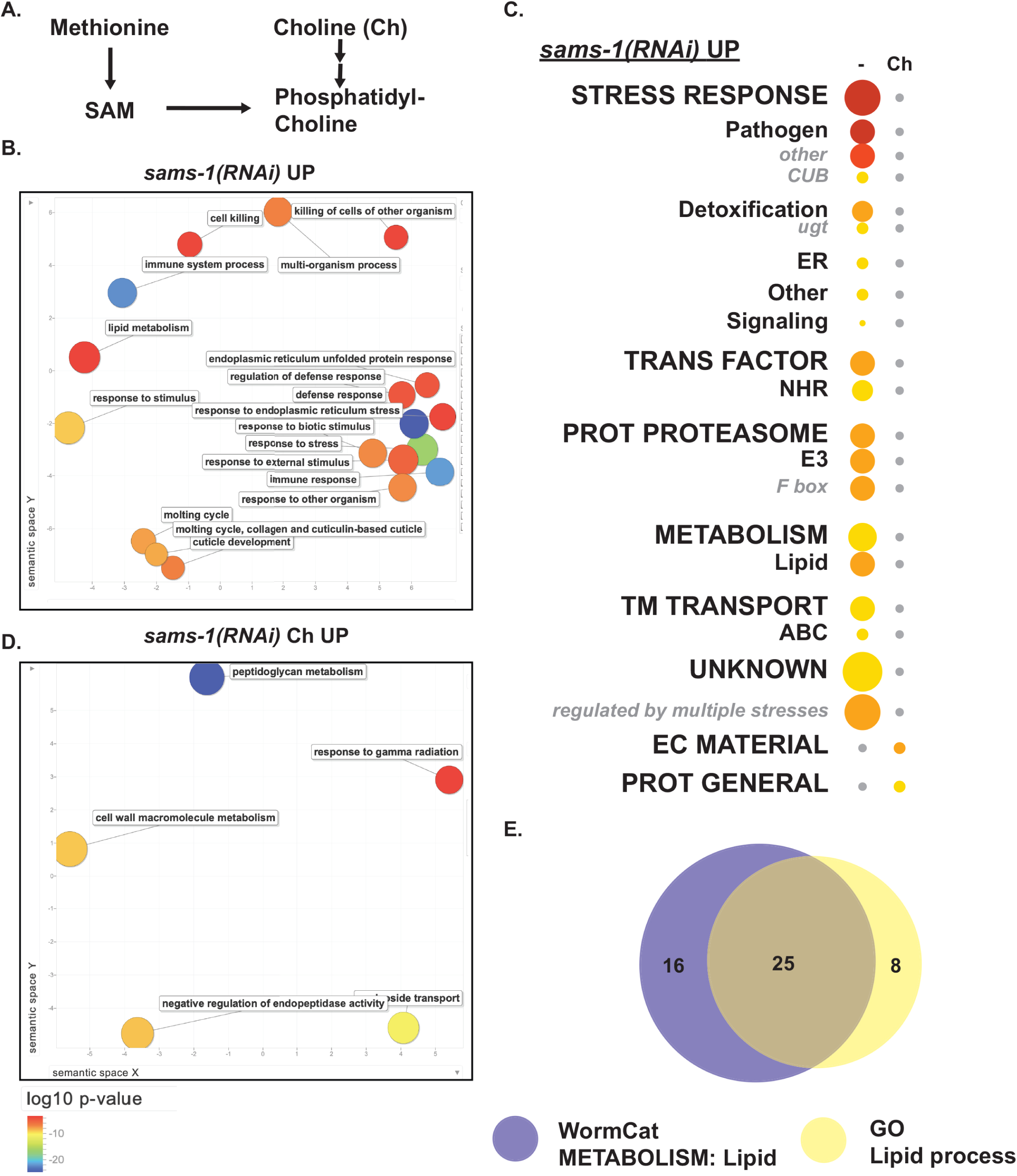
WormCat verifies known category enrichments *sams-1(RNAi)* upregulated genes. (**A**) Schematic showing metabolic pathways linking methionine, SAM, choline, and phosphatidylcholine. Gene expression microarray data for **B-D** were obtained from (Ding *et al*. 2015). (**B**) Semantic plot of GO enriched classifications generated by REVIGO (Supek *et al*. 2011) from *sams-1(RNAi)* Up genes. (**C**) WormCat visualization of categories enriched in genes upregulated in *sams-1(RNAi)* animals with and without choline supplementation in order of Cat1 strongest enrichment. Categories 2 and 3 are listed under each Category 1, with Category 2 or 3 sets that appeared independently of a Category 1 listed last. Bubble heat plot key is the same as Fig 1D. (**D**) *sams-1(RNAi)* Up plus choline (Ch) genes visualized by REVIGO. (**E**) Venn diagram showing overlap between WormCat *Metabolism: lipid* and GO *Lipid process* gene annotations. ABC, ATP-Binding Cassette; Ch, Choline; CUB, Complement C1r/C1s, Uegf, Bmp1 domain; EC Material, Extracellular Material; NHR, Nuclear Hormone Receptor; Prot General, Proteolysis General; Prot Proteasome, Proteolysis Proteasome; SAM, S-adenosylmethionine; TM Transport, Transmembrane Transport; ugt, UDP-glycosyltransferase

In order to identify GO terms enrichment with WormCat, we submitted genes up or down regulated 2-fold or more in *sams-1(RNAi)* animals to both WormCat and GOrilla (Eden *et al*. 2009). We used REVIGO (Supek *et al*. 2011) to visualize GO output. Both GOrilla/REVIGO (**Fig 2B**; **FigS2 A, B; Table S4**) and WormCat (**Fig 2C; Table S5**) identified categories of stress-response and metabolism linked to lipid accumulation in the genes that are upregulated upon *sams-1* RNAi, which is in agreement with our previous analysis (Ding *et al*. 2015). Interestingly, the relative importance of lipid metabolism is different in the two analyses. In the WormCat analysis, *Metabolism: lipid* was the third most enriched Cat2 category with a *P-*value of 1.2 x 10^−9^ (**Table S5**). In the GO analysis, however, *lipid metabolic process* was found with a modest enrichment of FDR corrected *P-*value = 5 x10^−2^ (**Table S4**). WormCat identified 41 genes in the *Metabolism: lipid* category, whereas GOrilla’s GO term search identified 33 genes in *lipid metabolic process* (**Fig 2E; Table S4**). Further inspection showed that six of the genes identified by solely by GOrilla were phospholipid lipases or phosphatases, one was an undefined hydrolase with no homology or domain similarity to genes with known lipid functions, and one was a transmembrane protein that may be better classified in other categories (see **Table S4** for GO lipid genes annotated by WormCat, tab 5 “GO_lipid_sams_up”). For example, lipases that hydrolyze phospholipids are the end points of metabolic pathways but produce second messengers acting in signaling pathways. One of these genes, Y69A2AL.2 has significant similarity to the human phospholipase A2 gene, PLA2G1B (BLAST *e* score of 2×10^−11^). This class of phospholipases cleave 3-sn-phosphoglycerides to produce the signaling molecule arachidonic acid (Xu *et al*. 2009); therefore a classification of *Signaling* is likely more reflective of its biological function than *Metabolism: lipid*. Taken together, WormCat identifies more genes that are directly relevant to the increased lipid storage phenotype observed with *sams-1(RNAi)* or *(lof)* animals (Walker *et al*. 2011; Smulan *et al*. 2016).

Next, we compared WormCat analysis of *sams-1(RNAi)* upregulated genes to the Gene Set Enrichment Analysis (GSEA) tool located in the WormBase suite (Angeles-Albores *et al*. 2016). GSEA, a GO based tool, identified similar categories as GOrilla with a concurrently high score for *lipid catabolic process* (**Fig S1)**. Our test set included 773 genes (**Table S5, tab4**); however, 286 of these genes were excluded from the GSEA analysis (**Table S6**), similar to the percentage excluded in a GOrilla analysis (Ding, et al. 2018). Unlike GOrilla, GSEA provides the user with gene IDs of excluded genes (**Table S6**). Therefore, we asked if these genes were excluded because their functions were undefined or if they were instead capable of classification. We found that 118 of the 286 excluded genes were classified as *Unknown* by WormCat (**Table S6**). However, 92 of the 476 genes GSEA included were also *Unknown* in WormCat analysis (**Table S5**, tab 4). Thus, the genes within this set that are classified as *Unknown* by WormCat only partially overlap with those that are excluded from GO analysis. Furthermore, WormCat classified 117 genes within the 286 genes excluded from GSEA, with 16 in non-coding categories and the remaining 101 in protein coding categories such as *Cytoskeleton*, *Metabolism* and *Proteolysis: proteasome* (**Table S6**). Thus, analysis of genes excluded from GO shows that an important fraction can be annotated and that *Unknown* WormCat categories are represented in both genes included and excluded from GO analysis.

Next, we used WormCat to analyze genes downregulated in *sams-1(RNAi)* animals. We noted enrichment in *Development: germline and mRNA function* categories in *sams-1(RNAi)* animals and that this enrichment is lost with choline treatment (**Fig S2D, Table S5).** This is consistent with the reduction in embryo production after *sams-1(RNAi)* and the rescue of fertility when PC levels are restored by choline supplementation (Walker *et al*. 2011; Ding *et al*. 2015). *Stress response* categories, however, are enriched in downregulated genes from both *sams-1(RNAi)* and *sams-1(RNAi)* choline treated animals (**Fig S2C; Table S5**). This appears to contrast with the complete loss of enrichment after choline treatment in the upregulated stress-response genes (**Fig 2C; Table S5**). However, inspection of the annotated gene lists returned by WormCat shows that the individual genes within the down-regulated *Stress response* category are different (**Fig S2E; Table S5**). Thus, on a gene by gene level, this data shows that the effects of choline supplementation are distinct for the up and downregulated genes in the *Stress response* category. In addition, this demonstrates that by providing both gene set enrichment and annotation of individual genes, WormCat provides a level of analysis that is difficult to achieve by traditional GO methods.

### Tau-tubulin kinases family are enriched in spermatogenic germlines

*C. elegans* is a robust model system for studying development and differentiation. Study of hermaphrodite germline development has been of particular interest, as it first produces sperm, after which it switches to oocyte production (Hubbard and Greenstein 2005). This concurs with distinct gene expression programs for both processes (Greenstein 2005; L’hernault 2006). Recently, the Kimble lab performed RNA-seq on dissected germlines from genetically female (*fog-2(q71))* and genetically male (*fem-3(q96*)) animals (Ortiz *et al*. 2014) (**Fig 3A**). Genes that were expressed in both germlines were called gender-neutral (GN), in contrast to genes that are specific to female (Oo, oogenic) or male (Sp, spermatogenic) germlines (Ortiz *et al*. 2014). We used WormCat to analyze the categories that were enriched in each dataset. We found that GN genes are strongly enriched for growth, DNA, transcription, and mRNA functions (**Fig 3B; Table S7**), which is expected because the germline is undergoing extensive mitotic and meiotic divisions. We further found that *Chromosome dynamics* and *Meiotic functions* were enriched in the GN dataset (**Fig 3C; Table S7**), as were *mRNA functions* of *Processing* and *Binding* (**Fig 3D; Table S7**). Oo genes were enriched for mRNA binding proteins, especially the zinc finger (ZF) class (**Fig 3D; Table S7**). These include such as maternally deposited *oma-1*, *pie-1*, *pos-1*, and *mex-1, mex-5* and *mex-6* mRNAs, which are known to function in oocytes (Lee and Schedl 2006) (**Table S7**). ZF proteins with unknown nucleic acid binding specificity were also enriched in the Oo dataset (**Fig 3D; Table S7**), suggesting that many of these may also be produced in the maternal germline. In an independent data set comparing RNA from germline-less (*glp-4(bn2))*, oocyte (*fem-3(gof))* and sperm-producing (*fem-1(lof))* animals by microarray analysis (Reinke *et al*. 2000), we also observed that categories in mRNA functions, transcription, development and cell cycle control were enriched (**Fig S3A-D, Table S8**).

**Figure 3:**
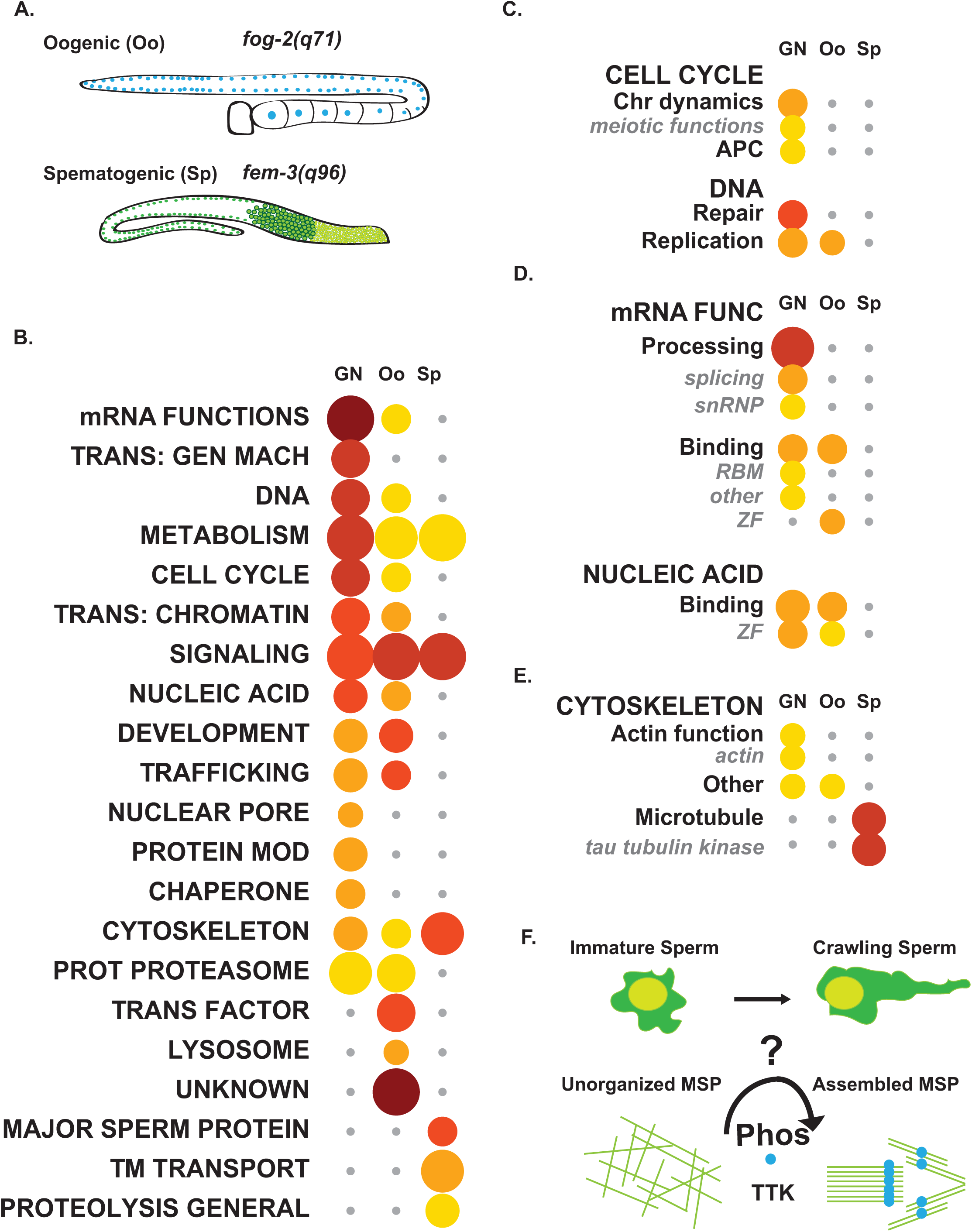
Analysis of germline-specific RNA-seq data identifies the tau tubulin kinase family as a male-specific category. (**A**) Schematic showing germlines used for female (top) or male (bottom)-specific RNA-seq analysis from Ortiz *et al*., and the mutant alleles to cause these phenotypes. (**B**) WormCat Category 1 analysis of Germline neutral (GN), Oogenic (Oo) or Spermatogenic (Sp) datasets ordered by most enriched in GN data. (**C-E**) Breakdown of WormCat enrichment from the Category 1 level for Cell Cycle (**C**), mRNA Functions and Nucleic Acid (**D**), and Cytoskeleton (**E**). Bubble heat plot key is the same as Fig 1D. (**F**) Schematic showing predicted phosphorylation and organization of MSPs during *C. elegans* sperm maturation based on WormCat findings. APC, Anaphase Promoting Complex; Chr Dynamics, Chromosome Dynamics; mRNA Func., mRNA Function; MSP, Major Sperm Protein; Phos, Phosphorylation; Protein Mod, Protein Modification; Prot Proteasome, Proteolysis Proteasome; RBM, RNA Binding Motif; TTK, Tau Tubulin Kinase; TM Transport, Transmembrane Transport; Trans: Gen Mach, Trans: Chromatin, Transcription: Chromatin; Transcription: General Machinery; Trans Factor, Transcription Factor; ZF, Zinc Finger

As expected, Sp genes are enriched for *Major Sperm Proteins* (MSPs), which are necessary for sperm crawling (**Fig 3B; Table S7**). Interestingly, a class of potential cytoskeletal regulators, *tau-tubulin kinases* (TTKs), were also enriched in Sp genes (64 of 71, *P-*value of 8.8 x10^−34^) (**Fig 3E; Table S7**). One TTK, *spe-6*, was previously isolated in a screen for spermatogenesis defects and is thought to be involved in phosphorylation of MSPs to allow the sperm to crawl (Varkey *et al*. 1993). Underscoring the potential importance of the TTKs in the male germline, WormCat also produced an enrichment in *tau tubulin kinases* in the Reinke, et al. spermatogenic gene sets (**Fig S3E, Table S8**). Thus, WormCat has identified a class of kinases that may be important for sperm-specific functions (**Fig 3F**).

To directly compare gene set enrichment from WormCat and GO, we analyzed each of these germline-enriched datasets with GOrilla and used REVIGO (Supek *et al*. 2011) for visualization (**Fig S4A-C, Fig S5A-B; Table S7, S8**). For the GN genes, the top 5 of the 544 significantly enriched categories were nucleic acid metabolic process (GO:0090304), nucleobase-containing compound metabolic process (GO:0006139), heterocycle metabolic process (GO:0046483), cellular aromatic compound metabolic process (GO:0006725), and organic cyclic compound metabolic process (GO:1901360) (**FigS4A, Table S7, see tabs 7, 8**). These GO categories are highly overlapping and are linked to multiple general processes involving nucleic acids. One gene GO:0006139, *gut-2*, an LSM RNA binding protein, was present in 23 different GO categories (**Table S7).** Comparison of these GO categories found that each contains genes placed in distinct WormCat categories. For example, *gut-2* was placed in *mRNA Functions* in WormCat, *ama-1*, the RNA Pol II large subunit, placed in *Transcription: General Machinery*, *brc-1*, the BRCA1 ortholog, placed in *DNA* and *nsun-5*, a mitochondrial RNA methyltransferase placed in *Metabolism: mitochondria*. These WormCat categories are the top five identified in the GN dataset (**Fig 3B, Table S7**). Thus, while WormCat and GO are both identify nucleic acid-related processed as among the most highly enriched in the GN dataset, the WormCat data is more concise and easily aligned with the molecular processes.

Within the spermatogenic datasets from Ortiz *et al*. and Reinke *et al*., WormCat identified a class of kinases, tau tubulin kinases (TTKs), that have the potential to function in sperm motility. General categories of phosphorus metabolic process (GO:0006793), phosphate-containing compound metabolic process (GO:0006796) and peptidyl-threonine phosphorylation (GO:0018107) were among the top five most enriched categories by GO from the Spermatogenic dataset, however, the TTKs as a group were not selectively identified from these very broad signaling categories in either spermatogenic data set (**Table S7, Table S8**). Thus, WormCat provided advantages over GO in the germline data sets by providing less redundant and more easily interpreted data and, most importantly, by identifying novel categories with potential links to biological function.

### Identification of post-embryonic tissue-specific gene expression categories

Improved technologies for cell-type-specific marker expression, nematode disruption, and deep sequencing of small RNA quantities have allowed construction of gene expression datasets from larval (Spencer *et al*. 2011) and adult somatic tissues (Kaletsky *et al*. 2018). To generate data from larval cell types, the Miller lab used cell-type specific tagged green fluorescent proteins to label a wide variety of larval tissues and examined mRNA expression in tiling microarrays (Spencer *et al*. 2011). RNA from each cell type would include tissue-specific, broadly expressed and ubiquitously expressed genes. To define cell-type specific transcripts, Spencer *et al*. designated *selectively enriched genes* as expressed more than 2-fold vs. the whole animal and as present in few cell types (Spencer *et al*. 2011). First, we performed WormCat analysis on the *selectively enriched* gene sets and found distinct gene set enrichments for each tissue type (**Fig 4A, Table S9**). For instance, body wall muscle (BWM) was enriched for *Muscle Function* and *Cytoskeleton* (**Fig 4B; Table S9**). The category *Metabolism* was enriched in both intestine (Int) and hypodermis (Hyp), whereas *Stress responses* appeared more specific for the intestine, and *Extracellular material* for the hypodermis (**Fig 4B, C; Table S9**). This likely reflects the role of the intestine in mediating contact with the bacterial diet after ingestion and the importance of the hypodermis for cuticle formation in larval development. While metabolic genes are expected to be required across multiple cell types, some cell types have specialized metabolic requirements. Both intestine and hypodermis are enriched for lipid metabolism genes at the Cat2 level. However, Cat3 analysis shows that sterol and sphingolipid genes drive this enrichment in the intestine while hypodermal lipid enrichment involves more broad categories with minor enrichments in *Metabolism: lipid: binding* and *Metabolism: lipid: lipase* (*P*-values of 4.51×10^−04^ and 2.86×10^−04^, which did not satisfy the FDR cutoff)(**Fig 4D; Table S9**). The Cat1 level analysis showed strong enrichment of transmembrane (TM) transporters in all tissues including the intestine, excretory cells and in neurons, however the Cat2 level shows enrichment of distinct classes of transporters (**Fig 4B; Table S9**) aligning with functions such as nutrient uptake, waste processing, and channel activity in each of these cell types.

**Figure 4:**
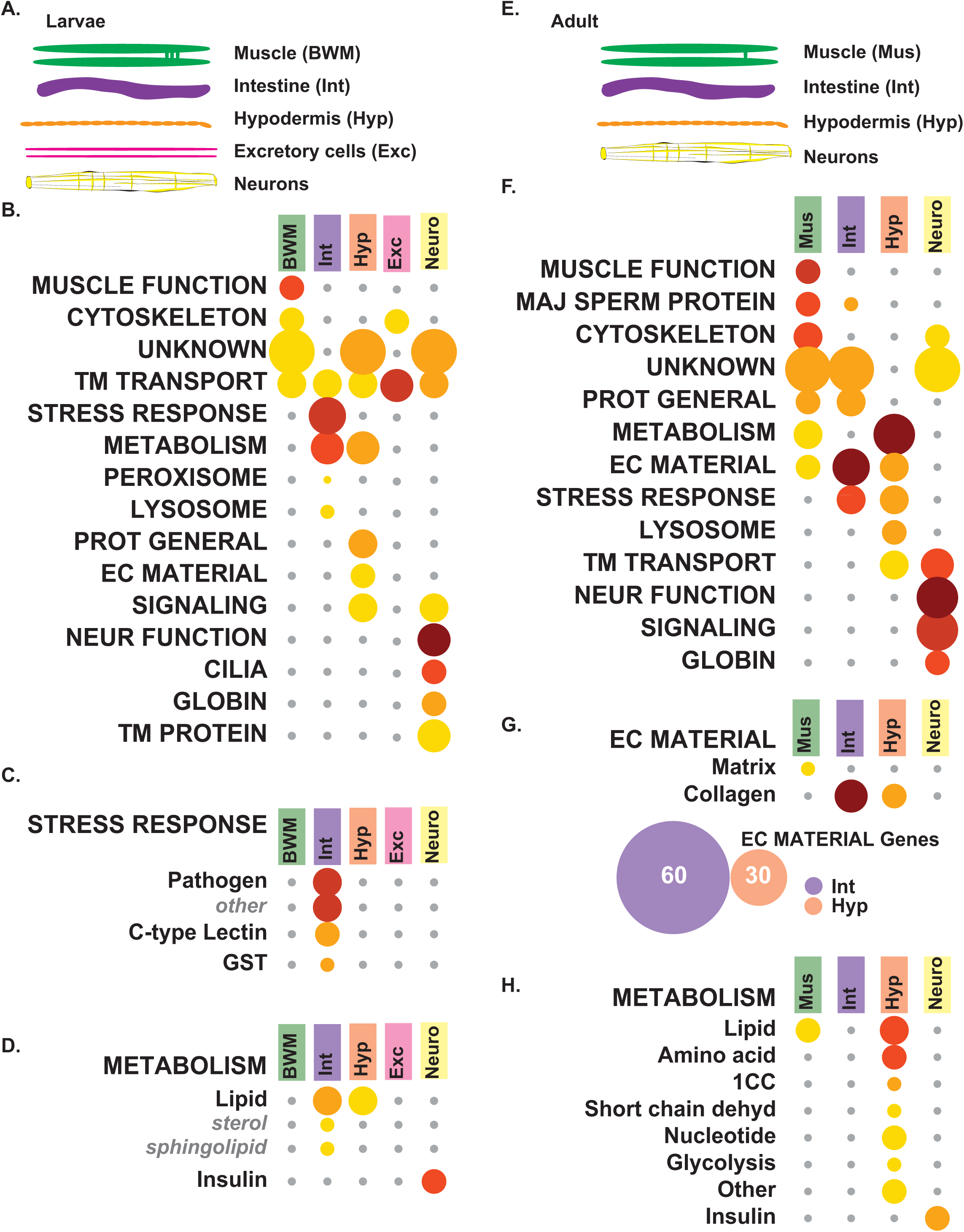
WormCat analysis of tissue-specific gene sets reveals the importance of the intestine in stress-responsive categories. (**A**) Diagram showing larval tissues isolated in tiling array data used in figures **B-D** from Spencer *et al*. (**B**) WormCat Category 1 enrichment for larval tissue-specific *selective enriched* gene sets shows differentiation of Body wall muscle (BWM), Intestine (Int), Hypodermis (Hyp), Excretory cells (Exe) and Neurons (Neuro). (**C-D**) Category 2 and 3 breakdown of Stress Response (**C**) and Metabolism (**D**). (**E**) Schematic showing adult tissues isolated for RNA-seq used in figures **F-I** from Kaletsky *et al*. (**F**) Category 1 analysis of enriched genes shows the differentiation of muscle and neuronal functions. (**G-H**) Category 2 and 3 breakdown of Extracellular Material gene enrichment including a Venn Diagram showing relationships between collagen genes in intestine and hypodermis (**G**), and Metabolism (**H**). Bubble heat plot key is the same as Fig 1D. 1CC, 1-Carbon Cycle; EC Material, Extracellular Material; GST, Glutathione-S-transferase; Maj Sperm Protein, Major Sperm Protein; Neur Function, Neuronal Function; Prot General, Proteolysis General; Short Chain Dehyd, Short Chain Dehydrogenase; TM Transport, Transmembrane Transport

Next we examined the data from Kaletsky *et al*., who performed RNA-seq from adult *C. elegans* sorted for muscle (Mus), intestinal (Int), hypodermal (Hyp) and neurons (Kaletsky *et al*. 2018) (**Fig 4E; Table S10**). They computationally separated genes to distinguish expression specificity, demarking “enriched”, “unique” and “ubiquitously” expressed categories. We used the “enriched” gene sets in WormCat analysis and found that WormCat correctly mapped muscle or neuronal genes to those cell types (**Fig 4F; Table S10**). At the Cat1 level, *Extracellular material* was enriched in muscle, hypodermis and intestine (**Fig 4F; Table S10**). At the Cat2 levels, *Extracellular material* diverged with *matrix* showing enrichment in muscle and *collagen* showing enrichment in intestine and hypodermis (**Fig 4G; Table S10**). However, the collagen genes enriched in intestine and hypodermis were distinct (**Fig 4G; Table S10**), perhaps reflecting differing roles for these collagens in the cuticle vs. in basement membranes. Distinguishing individual genes for this comparison is very cumbersome in commonly used GO servers and therefore represents an advantage of using WormCat. Previous studies found that two intestinal basement membrane collagens were produced in non-hypodermal tissues (Graham *et al*. 1997); however, this data suggests that others could be locally produced by the intestine. Kaletsky *et al*. also noted enrichment of metabolic function in adult hypodermis with GO analysis. Metabolic gene enrichment was also detected by WormCat analysis of their data (**Fig 4H; Table S10**), as well as in the larval data from Spencer *et al*. (**Fig 4D; Table S9**).

In our annotation strategy, we chose to restrict genes in categories such as *Neuronal function* to those that are specific to that tissue, and that have a described physiological function. Genes which functioned in neurons as well as other tissues were placed in more general molecular function-based categories. With this approach, we hoped to reduce false-positive identification of neuronal categories outside the nervous system, yet permit the identification of related, yet functionally less-specific groups. For example, while the WormCat analysis of the neuronal tissues in the Spencer *et al*. and Kaletsky *et al*. datasets showed strong enrichment of neuronal-specific categories, it also included categories of genes likely to function in both neurons and other tissues, or that contained genes that had not yet been classified *in vivo*. These categories include *Metabolism: insulin* (**Fig 4D, H; Table S10**), *Transmembrane (TM) transport, Signaling* (**Fig 4B, F; Table S10**) and *Transmembrane protein* (**Fig 4B; Table S10**). This is in line with the analysis by both Kaletsky *et al*. and Ritter et al. (Ritter *et al*. 2013)which also noted insulin expression across tissues and noted that more insulin genes were expressed at higher levels in adult neurons.

In order to distinguish the utility of WormCat from GO for the tissue-specific Spencer *et al*. and Kaletsky *et al*. datasets, we used GOrilla (Eden *et al*. 2009) to generate GO analysis and visualized the data with REVIGO (Supek *et al*. 2011) (**Figure S6-S8; Table S9, S10**). There were many similarities among the categories. For example, categories linked to the *Cytoskeleton* are highly enriched in the muscle datasets from Kaletsky *et al*. by GOrilla and WormCat (**Fig 4F, Fig S7A, Table S10**). In another example, *Stress response* categories were highly enriched by both WormCat and GO in the larval (Spencer *et al*. 2011) and adult (Murphy *et al*. 2003) intestine (**Fig 4F, Fig S6B, S7B, Table S10**). However, as shown above, WormCat identified the insulin gene family as strongly enriched in both the larval (**Fig 4D**) and adult (**Fig 4H**) neuronal tissue. Insulins were not identified as a class by our GO analysis. Instead, they were distributed among less specific categories such as biological regulation (GO:0065007), regulation of biological process (GO:0050789) and regulation of cellular process (GO:0050794) (**Fig S5, S6; Table S9, S10**). Thus, WormCat finds the major categories shown by GOrilla in the tissue-specific data and also identifies additional enriched groups.

The seven transmembrane protein family in *C. elegans* presented an annotation challenge. This class comprises around 8% of all protein-coding genes that seem likely to function in neurons, yet whose functions are undescribed (Robertson and Thomas 2006). Some have significant homology to mammalian G protein-coupled receptors (GPCRs), while others are nematode or *C. elegans* specific (Robertson and Thomas 2006). In order to identify and classify these proteins as accurately as possible, GPCRs with strong evidence for neuron-specific activity were placed in *Neuronal function*, while all other potential GPCRs were classified by protein domain and homology. For developing a list of potential GPCRs, we selected genes identified in WormBase as containing a transmembrane domain as well as those we initially annotated as GPCRs in the *Signaling* category. To recover any genes missed by these approaches, we added all *Unknown* proteins from our annotation list. We submitted the protein sequences for these genes to the NCBI Conserved Domain search tool (Marchler-Bauer *et al*. 2017) and selected all the genes in these groups that contained a seven-transmembrane (7TM) domain (**Fig 5A**). Next, we used BLASTP to determine the degree of homology to human GPCRs, which would reflect the conservation of function. Genes that had BLASTP scores of *e* < 0.05 on the NCBI server were classified in *Signaling: heteromeric G protein: receptor*. Those with *e* scores > 0.05 were classified as *TM protein: 7TM*, with class designated by WormBase in Cat3. Thus, genes with classified within *Neuronal function* or *Signaling* have a strong likelihood of GPCR function, whereas those in *TM protein: 7TM* have not been sufficiently defined. *Signaling: G protein* categories are enriched in neuronal genes sets from both Kaletsky *et al*. and Spencer *et al*. (**Fig 5B, C; Table S9, S10**) and 7TM proteins show enrichment in the larval pan-neuronal, *glr-1*-expressing neurons and motor neurons **(Fig 5C; Table S9, S10**). Thus, our annotation strategy allows separation of GPCRs with a highly likelihood of neuronal function, yet still permits enrichment of the larger class of 7TM proteins in neuronal tissues.

**Figure 5:**
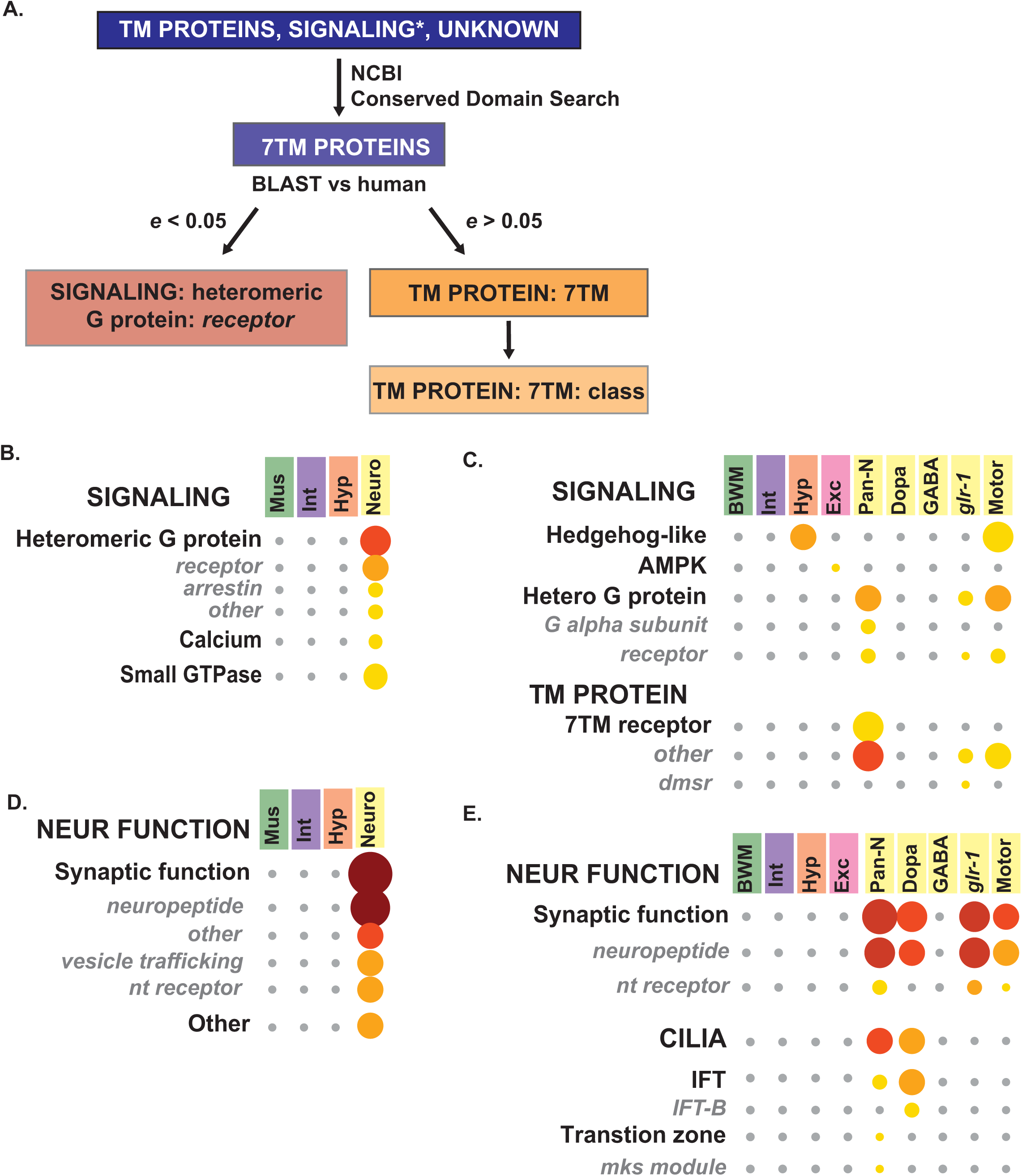
Detailed analysis of neuronal tissue-specific gene sets reveals specific enrichment for cilia gene expression on dopaminergic neurons. (**A**) Flow chart showing the process for annotating seven transmembrane (7 TM) proteins. *e* value is the statistical score provided by the NCBI BLAST server. Asterisk on Signaling notes that only predicted GPCRs within this category were submitted to the NCBI conserved domain server. (**B-E**) Breakdown of Neuronal Function to Category 2 and 3 from larval data in Kaletsky *et al*. (**B, D**) or adult data in Spencer *et al*. (**C, E**). 7TM receptor, Seven Transmembrane Receptor; BWM, Body Wall Muscle; dmsr, DroMyoSuppressin Receptor Related; Dopa, Dopaminergic Neurons; Exe, Excretory Cells; GABA, Gamma-Aminobutyric Acid-Specific Neurons; *glr-1*, Glutamate Receptor-Specific Neurons; Hetero G protein, Heterotrimeric G Protein; Hyp, Hypodermis; IFT, Intraflagellar Transport; Int, Intestine; mks module, Meckel-Gruber syndrome Module; Motor, Motor Neurons; nt Receptor, Neurotransmitter Receptor; Neuro, Neurons; Pan-N, Pan-Neuronal

In order to directly compare WormCat and GO on the larval neuronal data sets, we examined category enrichment of Spencer *et al*. pan-neuronal and motor neuron genes in GO by GOrilla (Eden *et al*. 2009), using REVIGO (Supek *et al*. 2011) for visualization (**Fig S6, S8; Table S9**). The most enriched category in the pan-neuronal or motor neuron datasets was G protein-coupled receptor signaling (GO:0007186). Next, we used WormCat to determine how we had annotated genes within GO:0007186 and found that this GO category included genes we had classified in *Signaling: Heteromeric G protein (G-alpha subunits and receptors)*, *Neuronal Function: Synaptic function* (neuropeptides and neurotransmitter receptors) and *TM protein: 7TM receptor* (**Fig 5C, Table S9**). While inclusion of the G protein signaling apparatus and neuropeptide ligands is appropriate for the broad category of G protein signaling, the GO categories do not differentiate between GPCRs with a high likelihood of function from the 7TM proteins that have not been functionally characterized. In addition, many of the *nlp* genes listed in GO:0007186 have not been functionally characterized and thus, it is not clear if they are bona fide GPCR ligands or could interact with other receptors outside of GPCR signaling (Li and Kim 2008). Therefore, WormCat improves on GO analysis for these datasets by providing more nuanced information on the function of these genes in GPCR pathways.

Neuronal genes from adult (Kaletsky *et al*. 2018) and larval gene sets (Spencer *et al*. 2011) also showed strong enrichment in Cat2 and Cat3 classifications within *Neuronal function*, such as *Synaptic function*, *neuropeptide*, and *neurotransmitter (nt) receptor* (**Fig 5D, E; Tables S9-S10**). *Cilia* genes were also enriched in the pan-neuronal and dopaminergic larval gene sets (**Fig 5D; Table S9**). Neurons are the only ciliated cells in *C. elegans* and cilia occur on multiple neuronal subtypes (Inglis *et al*. 2007). However, all dopaminergic neurons are ciliated (Inglis *et al*. 2007), and are therefore more likely to show enrichment. Taken together, our WormCat analysis of these large tissue-specific gene sets provides a detailed view of gene classes specific to muscle, hypodermis, intestine, and neurons in larvae and adults. We have identified differential enrichment in lipid metabolism genes and collagens from intestine and hypodermis, defined a classification system for GPCRs and 7TMs and identified *Cilia* as a major enriched category in dopaminergic neurons. Much of this information goes beyond what is revealed in GO analysis and provides predictions that can be useful to design future studies. Identification of these types of nuanced tissue-specific patterns is an important step to understanding how specific cell types function.

### Drug interactions limiting lifespan induce changes in sterol metabolism

*C. elegans* is particularly suited to studies determining gene expression changes in response to a panel of treatments in a whole animal, and to correlate these changes to physiological function. For example, Admasu *et al*. generated a complex gene expression dataset by performing parallel RNA-seq on animals treated with five lifespan-increasing drugs that affect distinct pathways (Allantoin, Rapamycin, Metformin, Psora-5, and Rifampicin). They used five pairwise combinations and three triple drug combinations to determine if any combination lead to further lifespan extension, and to identify gene expression profiles associated with increased longevity (Admasu *et al*. 2018). They found that one triple drug combination (Rifa/Psora/Allan) activated lipogenic metabolism through the transcription factor SBP*-*1/SREBP*-*1 and determined that the drug-induced longevity was dependent on SBP-1 function (Admasu *et al*. 2018). The authors also made the striking observation that a distinct triple drug combination (Rifa/Rapa/Psora) reduced lifespan, even though each single drug or drug pairs increased longevity (Admasu *et al*. 2018). To determine if any gene expression categories might explain this effect, we used WormCat to analyze category enrichment for the up and downregulated genes for each single drug, pairwise or triple drug combination (**Fig 6A, Figs S9, S10; Tables S11, S12**). Similar to the author’s KEGG analysis (Admasu *et al*. 2018), we observed *Metabolism: lipid* enrichment in long-lived Rifa/Rapa/Psora-treated animals (**Fig 6A, Table S11**), however, we also noted that *Metabolism: lipid* was enriched in all three combinations with WormCat. Next, we examined the up and downregulated genes to determine if any categories correlated with the failure to survive in the Rifa/Rapa/Psora treated animals. We did not find category signatures in the downregulated genes that appeared to correlate with the decrease in longevity (**Fig S10**; **Table S12**). However, upregulated genes from the short-lived Rifa/Rapa/Psora treated animals were enriched in another specific class of lipid metabolic genes: sterol metabolism (**Fig 6A, Fig S9**). Closer examination of the single and pairwise combinations showed that the enrichment of sterol metabolic genes only appeared in the triple combination with poor survival (**Fig 6B**). *C. elegans* do not use cholesterol as a membrane component (Ashrafi 2007). Thus, this category does not include cholesterol synthesis genes, but does include genes involved in modification of sterols, for example, in steroid hormone production (Watts and Ristow 2017). Examination of individual genes (**Table S11**, Tab 18 Sterol Genes) showed that five of the 19 had lifespan phenotypes and four had lethality related phenotypes in WormBase, consistent with their effects on survival in Admasu *et al*. Furthermore, Murphy *et al*. showed that three of the 19 sterol genes are upregulated in another long-lived model, *daf-2(mu150)*, and two of these, *stdh-1* and *stdh-3* are required for lifespan extension in *daf-2(mu150)* animals (Murphy *et al*. 2003). Thus, the category enrichments captured by WormCat for this drug study have identified sterol metabolism genes as potential players in the paradoxical lifespan shortening effects of the Rifa/Rapa/Psora combination.

**Figure 6:**
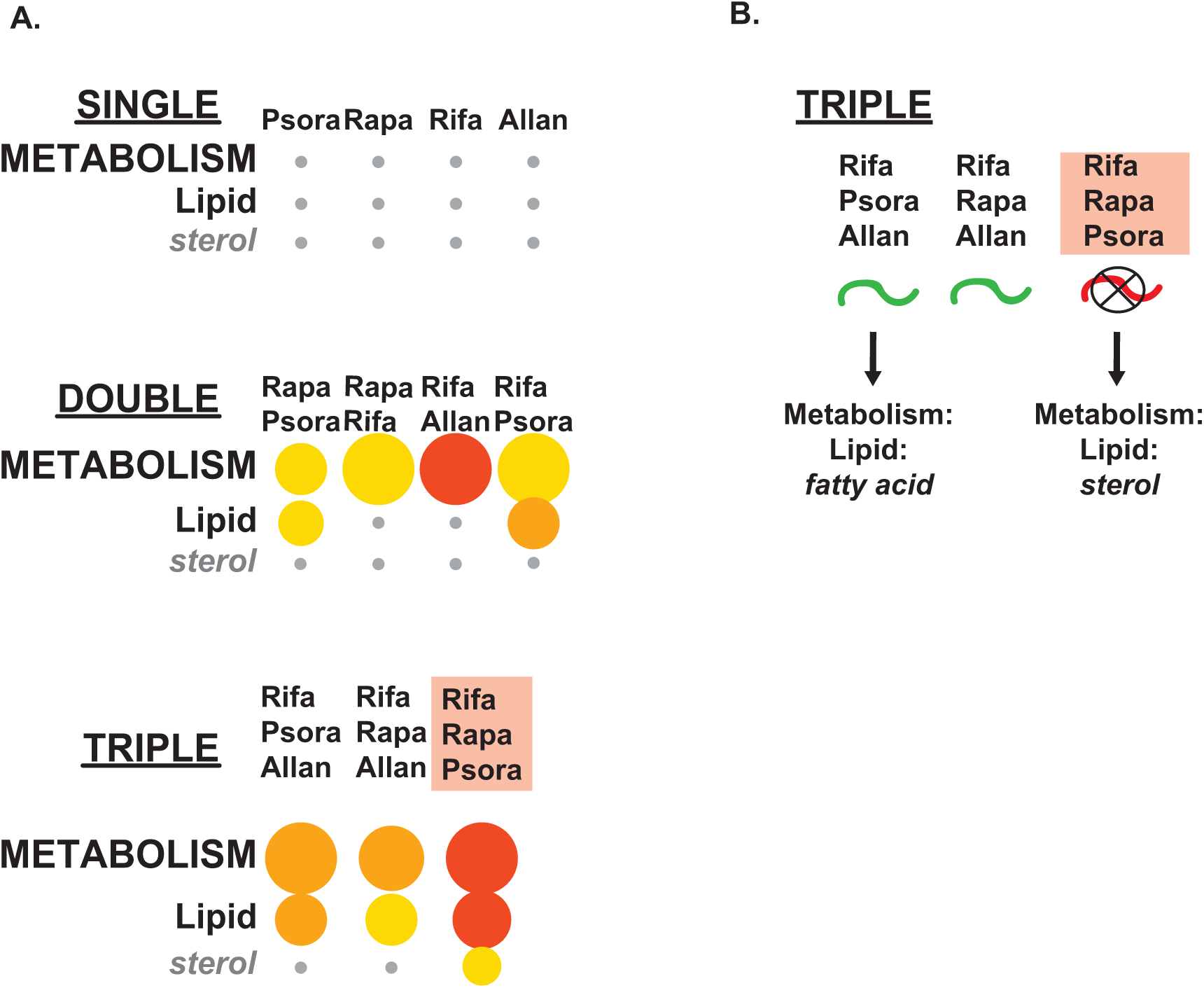
WormCat analysis of RNA-seq data from *C. elegans* treated with combinations of lifespan-lengthening drugs reveals the emergence of sterol metabolism in drug combinations limiting survival. (**A**) Comparison of *Metabolism: lipid: sterol* enrichment in single, double and triple-drug combinations shows sterol emergence in the Rifa/Rapa/Psora gene set (Admasu *et al*. 2018). (**B**) Diagram showing a summary of data from lifespan changes after triple drug treatment from Admasu *et al*. Pink box denotes drug combination that causes premature death. Bubble heat plot key is the same as Fig 1D. Allan, Allantoin; Psora, Psora-4; Rapa, Rapamycin; Rifa, Rifampicin

In order to compare gene set enrichment of the triple drug combinations from WormCat with GO, we analyzed upregulated genes from the Rifa/Psora/Allan, Rifa/Rapa/Allan and Rifa/Rapa/Psora treated animals in GOrilla (Eden, 2009) and visualized the data with REVIGO (Supek, 2011) (**Fig S11; Table S11**). WormCat and GO showed multiple similarities. For example, WormCat and GO identified extracellular matrix-linked categories in all three triple combinations (WormCat: EC MATERIAL; GOrilla: GO:0030198: extracellular matrix organization) (**Fig S9; Table S11**). However, WormCat identified *Metabolism: lipid* in all three combinations, whereas GO analysis by GOrilla only identified categories linked to lipid metabolism (GO:0006629: lipid metabolic process (*q* = 5.63×10^−03^), GO:0044255 cellular lipid metabolic process (*q* = 1.49×10^−02^) and GO:0006631 fatty acid metabolic process (*q* = 2.16×10^−02^)) in the Rifa/Rapa/Psora dataset (**Table S11**). WormCat also showed a much higher enrichment score for *Metabolism: lipid*, *p* = 2.00 x10^−14^) (**Table S11**). Thus, as in the *sams-1* microarray data discussed previously, WormCat provides an improved tool for determining enrichment of metabolic genes.

WormCat also found an enrichment of transcription factors in each of the triple combinations, with specific enrichments in nuclear hormone receptors and homeodomain genes in the Rifa/Psora/Allan upregulated set (**Fig S9**) Enrichments of nuclear hormone receptors in *C. elegans* is potentially of interest as they may regulate multiple metabolic regulatory networks (Arda *et al*. 2010). However, GOrilla only identified categories linked to transcription factors (GO:0006355: regulation of transcription, DNA-templated, GO:0051252: regulation of RNA metabolic process, GO:2001141: regulation of RNA biosynthetic process, GO:1903506 regulation of nucleic acid-templated transcription and GO:0019219 regulation of nucleobase-containing compound metabolic process) in the Rifa/Psora/Allan dataset. No individual class of transcription factors were identified in any of the triple combinations by GO (**Table S11**), thus WormCat offers a clear advantage over GO by providing increased coverage across diverse categories of gene function.

### Identification of gene set enrichments in RNAi screening data

In order to use WormCat to analyze genome-scale RNAi screening data, we mapped WormCat annotations to the list of genes in the Ahringer library (Kamath *et al*. 2003) (**Table S13**). To test this approach, we used data from the Roth lab who screened the Ahringer library for changes in glycogen storage in *C. elegans* and identified more than 600 genes, scored as glycogen high, glycogen low and abnormal localization (Lamacchia *et al*. 2015) (**Fig 7A, Table S14**). The authors functionally classified all hits from the screen with an in-house annotation list, graphed the percentage within each group, and noted high percentages of genes with roles in metabolism (electron transport chain), signaling, protein synthesis or stability, and trafficking (Lamacchia *et al*. 2015), however, they were unable to assign statistical significance to any of the groups. WormCat identified similar groups as the LaMacchia *et al*. functional classification for the glycogen low candidates. For example, we identified *Metabolism: mitochondria*, complex I, III, IV, and V and found that these categories were statistically enriched (**Fig 7B; Table S14**). However, signaling was not enriched (**Table S14**). Thus, WormCat is able to identify statistically relevant pathways in genome-scale RNAi screen data.

**Figure 7:**
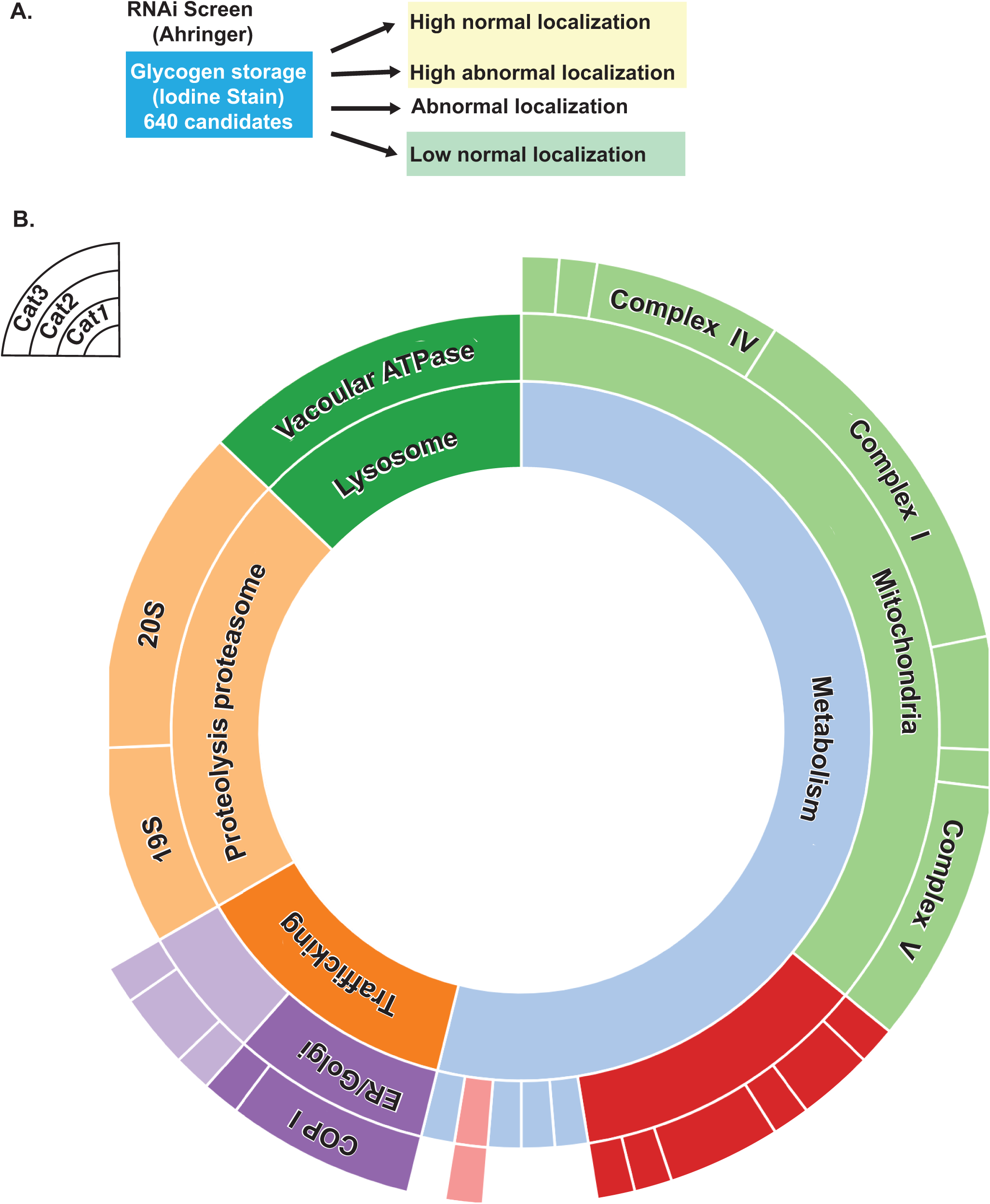
WormCat analysis of a genome-scale RNAi screen quantitates categories of candidate genes. (**A**) Schematic of the RNAi screen from LaMacchia *et al*. identifying candidate genes that altered glycogen staining. (**B**) Sunburst diagram from low glycogen candidates showing significantly enriched categories.

To provide a direct comparison between WormCat and GO with this data set, we determined the GO term associated with the glycogen low data by GOrilla (Eden *et al*. 2009) and visualized the data with REVIGO (Supek *et al*. 2011) (**Figure S12; Table S14**). 185 separate GO terms were identified in this data set compared to the four Cat1 level terms identified by WormCat (*Metabolism, Lysosome, Proteolysis Proteasome* and *Trafficking*) (**Fig 7B, Table S14**). WormCat also finds a limited number of Cat2 groupings within these sets including *Metabolism: mitochondria, Lysosome: vacuolar ATPase, Proteolysis Proteasome:19S, 20S*, and *Trafficking:ER/Golgi*) (**Fig 7B, Table S14**). This large difference in number of significantly enriched categories stems from the multiple, overlapping categories present in the GO analysis. For example, the mitochondrial gene *cyc-1* (Cytochrome C oxidase) is represented in 87 of the GO terms, whereas the annotation in WormCat is *METABOLISM: mitochondria* (**Table S14, tab 8**). Similarly, the vacuolar ATPase *vha-6* is represented in 39 of GO terms returned, the proteasomal component *psb-7* is present in 23, and the ER/Golgi COP I component Y71F9AL.17 is in 21 (see **Table S14**, tabs 9-11). This GO term redundancy provides the user with a complex, hard to interpret list. In addition, GO terms that are repeated fewer times (such as those containing the trafficking gene Y71F9AL.17) become marginalized in a complex list. Thus, with this dataset WormCat provides easily distinguished categories with clear links to biological or molecular function. The GO terms show the same genes repeated in a large fraction of the categories and obscure categories with less gene redundancy.

## Discussion

### WormCat provides new insights into comparative RNA-seq data

Current technology allows for the routine use of genome-scale experiments for the generation of gene expression data. The goal of these experiments is often to identify classes of genes that add insight to biological functions, as well as to highlight selected genes for individual analysis. GO analysis, while widely used, is difficult to apply to datasets with multiple combinations of treatments or genetic perturbations. Further, for *C. elegans*, current GO analysis is often inaccurate and misses useful physiological and molecular information. Here we have shown that WormCat can annotate gene categories, provide enrichment statistics, and display user-friendly graphics for gene sets identified from *C. elegans* gene expression studies. Furthermore, our visualization strategy allows comparison across multiple datasets, facilitating identification of categories that can be linked to shared biological functions.

Our initial, script-based, smaller-scale version of WormCat highlighted changes in metabolic gene expression in *C. elegans* with changes in levels of the methyl donor S-adenosylmethionine (SAM) or methyltransferases modifying H3K4me3 (Ding *et al*. 2018). In this study, we have expanded the annotation list, developed a web-based server, and added an additional graphical output. We used WormCat to successfully analyze data from metabolic, tissue-specific, and drug-induced expression changes. This analysis provides not only validation and use-case examples but also additional insights into the known gene expression patterns. For example, our examination of germline gene expression datasets from the Kimble and Kim labs (Reinke *et al*. 2000; Ortiz *et al*. 2014) identified a large class of microtubule kinases (tau tubulin kinases, TTK) as enriched in spermatogenic gene sets and as a co-enriched gene set with major sperm proteins (MSPs). One TTK, *spe-6*, has been previously identified in a screen for mutants with defects in sperm development (Varkey *et al*. 1993). Our results suggest that many genes in this family could have important functions in spermatogenesis and that the appearance of MSPs and TTKs in a dataset could also serve as a marker for maleness. Finally, we used WormCat to analyze a dataset consisting of RNA-seq from C*. elegans* treated with multiple lifespan changing drugs alone or in combination, plus one mutation animal strain that extends lifespan (Admasu *et al*. 2018). The classification and graphical output allowed us to identify upregulation of sterol metabolism genes in a triple-drug combination that was not present in the single or double drug treatments. Thus, WormCat identified a gene set that may be important for the effects of the lifespan-altering drugs in this assay.

### Strengths and weaknesses of WormCat

We developed WormCat to overcome some of the limitations of GO analysis when analyzing *C. elegans* gene expression data and to utilize specific phenotype data available in WormBase. In addition, we specifically engineered WormCat to classify data for identification of co-expressed or co-functioning gene sets. Finally, we developed two graphical outputs, a scaled heat map/bubble plot and a sunburst plot. The modular nature of the bubble plot allows multiple datasets to be grouped and compared, while the sunburst plot gives a concise view of single datasets, as may be obtained with screening data. Our validation with random gene testing and analysis of *C. elegans* gene expression data from metabolic, tissue-specific, and drug-treated animals shows that WormCat is a robust tool that provides biologically relevant gene enrichment information. There are three main areas that WormCat provides an advantage over using GO that are apparent in our case studies. First, as discussed above, we found that in some of our test cases, WormCat identified broader sets of genes within categories or categories that were not identified by GO. Second, the WormCat output is much easier to interpret; the bubble charts provide intuitive visualization and the tables provide clear access to the enrichment statistics and annotation of the input genes. Third, the availability of the annotations for each input gene enables comparisons between genes in categories. For example, we found that while *Extracellular material: collagen* was enriched in both intestine and hypoderm in the Kaletsky *et al*. data set, the genes were non-overlapping, suggesting tissue-specific expression of collagen genes. This comparison would be difficult to make with GO, as many common GO servers do not supply the genes with each category in an easily accessible manner. Directly comparing the genes within WormCat and GO categories from our previously published dataset of gene expression after *sams-1* knockdown, we found that WormCat identified a broader set of lipid metabolic genes than GO analysis from GOrilla and that the genes identified only by GO analysis might be better classified in different categories to reflect their biological functions. Thus, WormCat provides an alternative to GO with advantages in output that improve data interpretation and access to gene annotations that allow deeper comparisons among categories. In some cases, WormCat also identifies categories that are not found by GO.

However, there are several limitations to WormCat. First, while multiple researchers with varied expertise curated our annotation list, some genes may be mis-annotated, or some Cat2 or Cat3 groups may fit better in other Cat1 classifications. We will update the WormCat annotation list at periodic intervals while providing access to the previous annotation lists. Second, each *C. elegans* gene was given a single, nested annotation, rather than a group of annotations as in GO. We chose to prioritize the visualization of enriched gene sets in this instance, using a single annotation per gene to permit graphing in scaled heat maps. Access to the program and annotation lists for the local application also allows users to customize the annotation lists according to their preferences.

Annotation lists of genome-scale data are likely to contain errors. We have defined several sources of error and have taken corrective steps. In some cases, a gene may be simply mis-annotated. For example, a component of the *General transcription machinery* was placed in *Signaling* by the annotator. In others, the classification system may be incorrect. An example of this would be classifying enzymes that modify small molecules as protein modification. To estimate the mis-classification error rate, we generated a list of 3000 random WormBase IDs. We mapped each ID to our annotation list and rechecked the annotations. We found 29/2294 genes (1.3%) whose annotations were incorrect by our criteria (13 of these were Unknown genes which could be classified in other categories). This suggests around 300 genes in the entire dataset may be mis-annotated by our criteria, many representing Unknown genes which could acquire classification. We will periodically update the WormCat annotation lists to accommodate new gene information and correct errors.

It is important to note that some gene classifications depend on criteria that are open for interpretation. For example, transcription factors regulating genes within a pathway are grouped within a linked category to allow identification of co-functioning genes. For instance, *efl-1*, a master regulator of cell cycle genes is annotated as *Cell cycle: transcriptional regulator* instead of with the more broadly acting trans-regulatory factors in *Transcription factor: E2F*. To allow for different interpretations of the annotation strategy, we have set up a GitHub site (https://github.com/dphiggs01/wormcat) where the annotation list and scripts for executing WormCat can be downloaded and customized by the user to accommodate differences in annotation preference.

The value of gene set enrichment is also highly dependent on the criteria used to specify the regulated genes. In the present study, we used the same criteria as the respective authors, except that we separated up and downregulated genes where necessary. For example, in the Kaletsky *et al*. tissue-specific data, the authors provided data for all genes expressed in each tissue, enriched genes (expressed at FDR great than 0.05, and log_2_ fold change greater than 2 relative to other tissues), or unique genes (log_2_ RPKM greater than 5) significantly differentially expressed in comparison to the expression of each of the three other tissues (FDR greater than 0.05, log_2_ fold change greater than 2 for each comparison) (Kaletsky *et al*. 2018). We found the best resolution of WormCat categories between the tissues occurred with the enriched datasets, rather than with all genes or unique gene sets. This suggests that gene lists with all expressed genes may require more stringent statistical cutoffs, but also that WormCat may not be as suited to highly filtered data.

### Application to other organisms

By developing WormCat specifically for analyzing *C. elegans* gene sets, we were able to take advantage of available data on WormBase but limited the applicability of our annotation list with other organisms. Although researchers in mammalian fields can access pathway analysis pipelines such as Ingenuity Pathway Analysis (Qiagen, (Kramer *et al*. 2014)) that are focused on identifying functionally linked genes, these programs do not necessarily provide a simple graphical output for comparative analysis. WormCat analysis generating the scaled heat/bubble charts can be adapted for use with other organisms by running the program locally with altered annotation lists. Replacing gene IDs and the Cat1, Cat2 and Cat3 values with any annotation allows customization of the pipeline to any other organism. Thus, the modular nature of WormCat allows adaptation to multiple annotation strategies within *C. elegans* or to other organisms, allowing a streamlined visualization for examining genome-scale expression or screen data.

## Acknowledgments

We wish to thank members of the Walker and Walhout labs for helpful discussion. Funding to A.K.W NIH NIA 1R01AG053355. A.J.M.W. grants NIH grants DK068429 and GM122502.

