## Supplementary material for "WormCat: an online tool for annotation and visualization of *Caenorhabditis elegans* genome-scale data": Figure S1

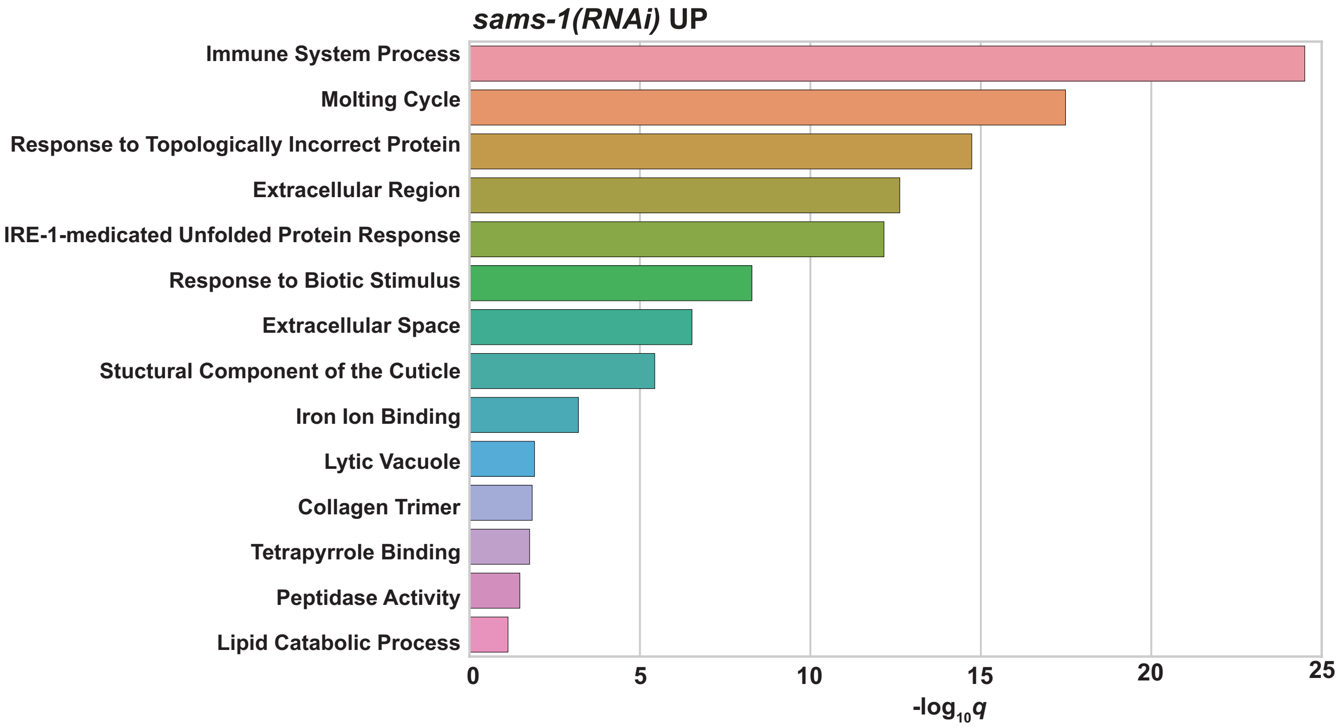


**Figure S1. GO Analysis of upregulated genes from *sams-1(RNAi)* animals by Gene Set Enrichment Analysis.** Bar graph showing GO categories returned from *sams-1(RNAi)* upregulated genes (Ding *et al.* 2015) by the WormBase Gene Set Enrichment Analysis tool (Angeles-Albores *et al.* 2016).
