## Supplementary material for "WormCat: an online tool for annotation and visualization of *Caenorhabditis elegans* genome-scale data": Figure S2

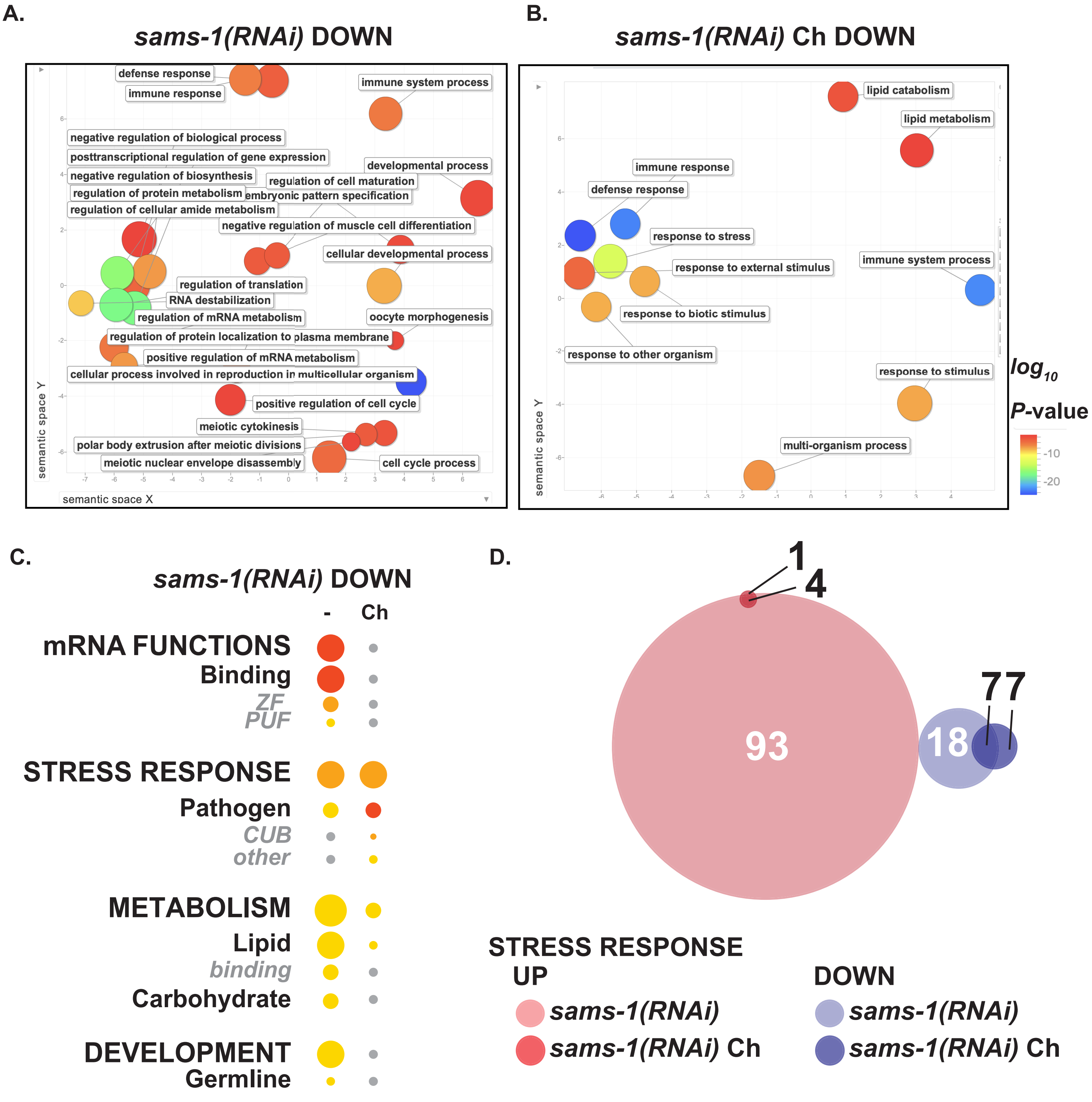


**Figure S2. WormCat verifies known category enrichments from *sams-1(RNAi)* downregulated genes.** (**A-B**) REGIVO semantic plots of *sams-1(RNAi)* downregulated genes from untreated (**A**) and choline (Ch) treated (**B**) animals. (**C**) WormCat bubble heat plots comparing *sams-1(RNAi)* with and without choline. Gene expression microarray data for **A-C** were obtained from Ding *et al*., 2015. Bubble heat plot key is the same as **Fig 1D**. CUB, Complement C1r/C1s, Uegf, Bmp1 Domain; PUF, Pumilio and *fem-3* mRNA Binding Factor; ZF, Zinc Finger. (**D**) Venn diagrams showing overlap between *Stress Response* genes in *sams-1(RNAi)* up (pink) or downregulated genes (blue).
