## Supplementary material for "WormCat: an online tool for annotation and visualization of *Caenorhabditis elegans* genome-scale data": Figure S3

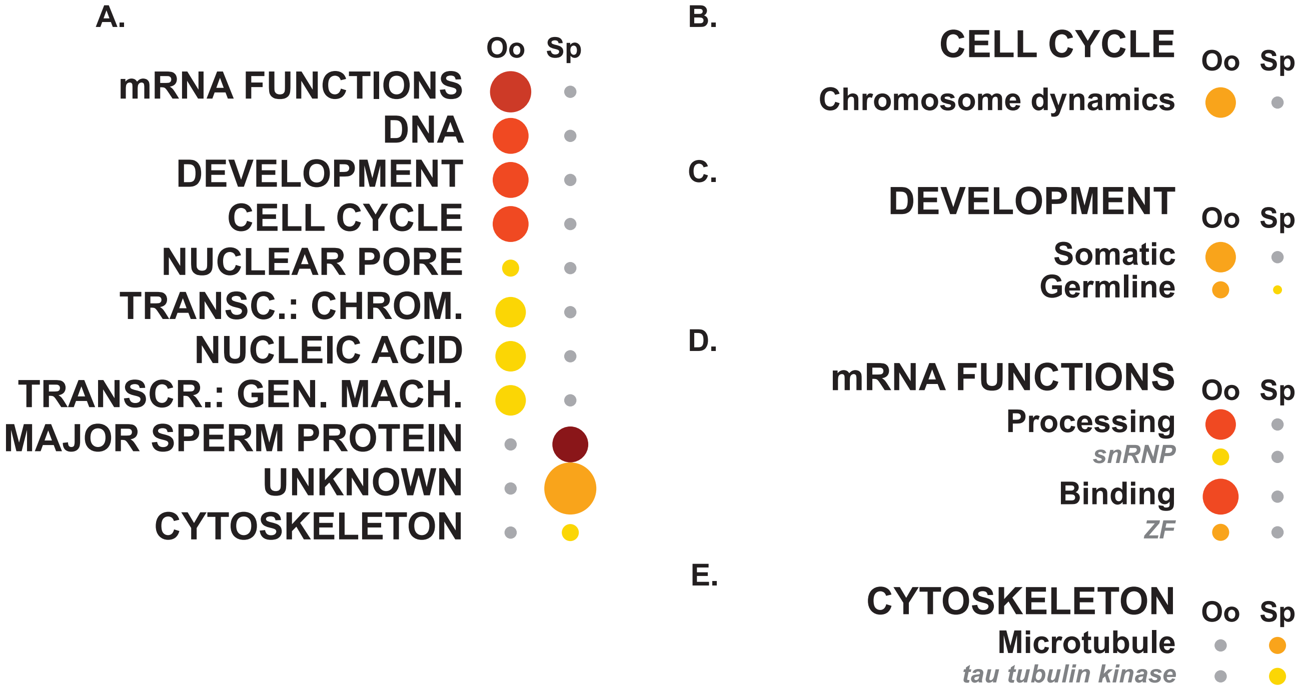


**Figure S3: WormCat analysis of germline-specific microarray data identifies the tau tubulin kinase family as a male-specific category**. (**A**) Category 1 analysis of Oogenic (Oo) or Spermatogenic (Sp) data sets ordered by most enriched in Oo data. Breakdown of data from the Category 1 level for Cell cycle (**B**), Development (**C**), mRNA Functions (**D**), or Cytoskeleton (**E**). All data is from Reinke et al. (Reinke et al., 2000). Gen. Trans. Machinery, General Transcription Machinery; Trans. Chromatin, Transcription: Chromatin; ZF, zinc finger.
