## Supplementary material for "WormCat: an online tool for annotation and visualization of *Caenorhabditis elegans* genome-scale data": Figure S4

**A.** **Germline Neutral (GN)**

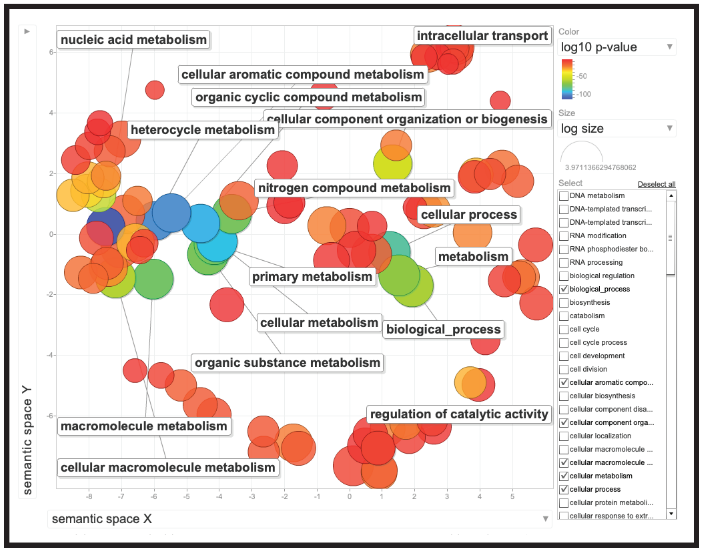


**B.** **Oogenic (Oo)**

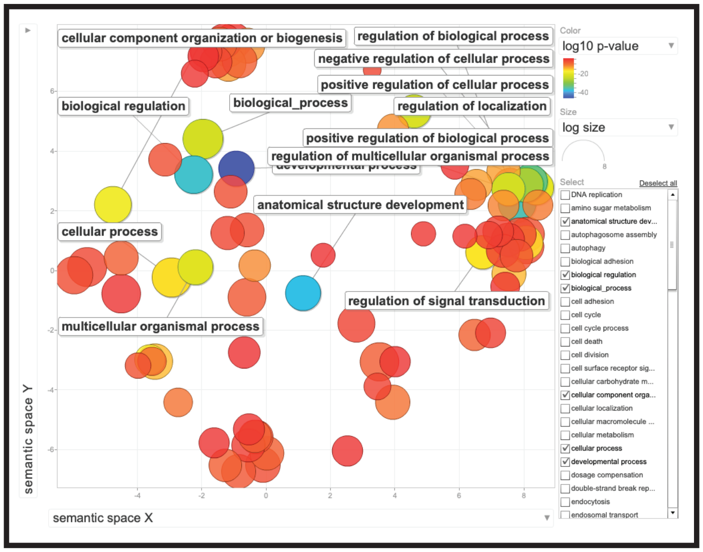


**C.** **Spermatogenic (Sp)**

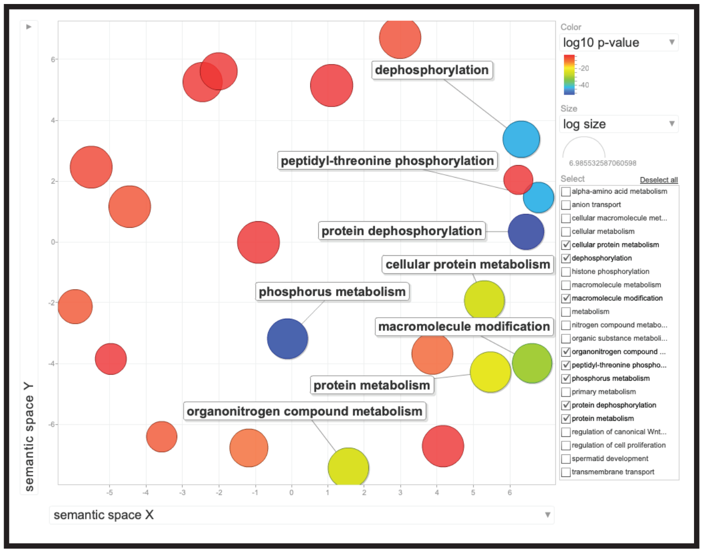


**Figure S4: GO analysis visualized by Revigo of germline RNA seq data from Ortiz, et al.** Sematic graphs of GO analysis generated by GOrilla (Eden *et al.* 2009) and visualized by Revigo (Supek *et al.* 2011) of Gender Neutral (**A**), Oogenic (**B**) and Spermatogenic (**C**) germlines (Ortiz *et al.* 2014).
