## Supplementary material for "WormCat: an online tool for annotation and visualization of *Caenorhabditis elegans* genome-scale data": Figure S5

**A.** **Oogenic (Oo)**

**B.** **Spermatogenic (Sp)**

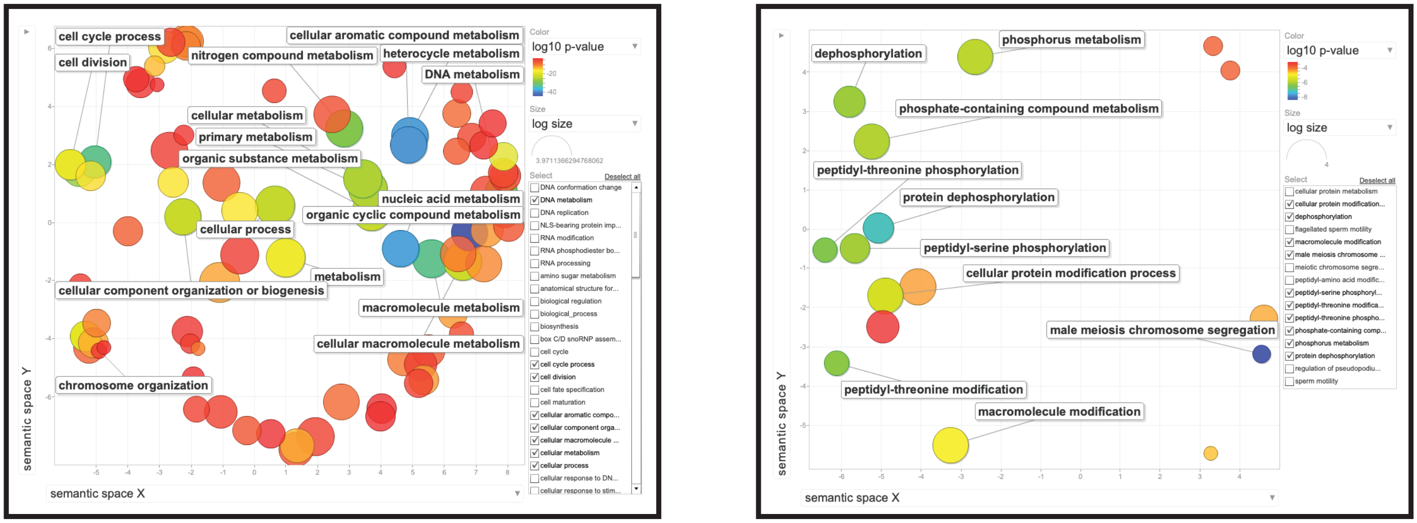


**Figure S5: GO analysis visualized by Revigo of germline microarray data from Reinke, et al.** Sematic graphs of GO analysis generated by GOrilla (Eden *et al.* 2009) and visualized by Revigo (Supek *et al.* 2011) of Oogenic (**A**) and Spermatogenic (**B**) germlines (Reinke *et al.* 2000).
