## Supplementary material for "WormCat: an online tool for annotation and visualization of *Caenorhabditis elegans* genome-scale data": Figure S6

Holdorf et al. Figure S6

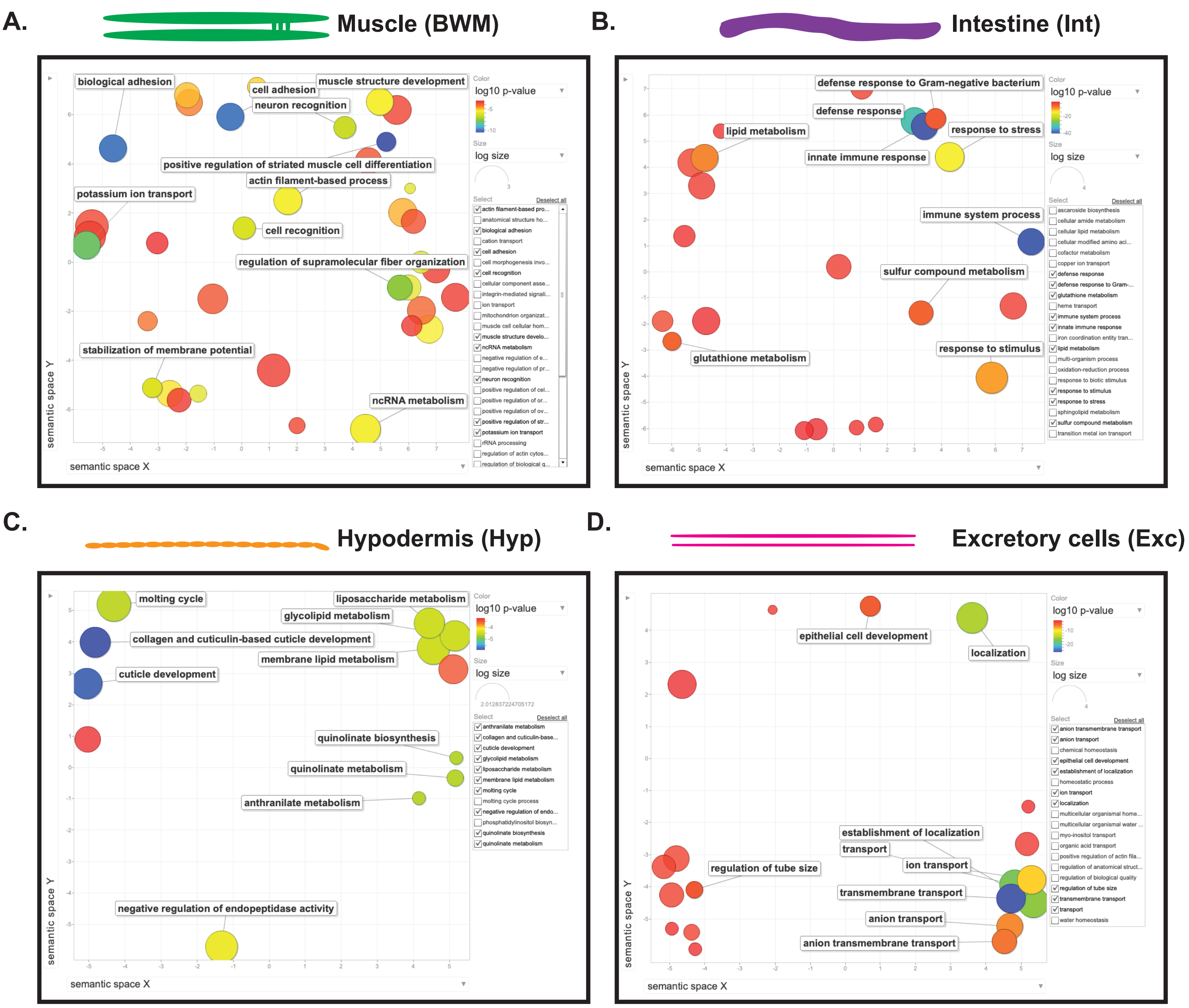


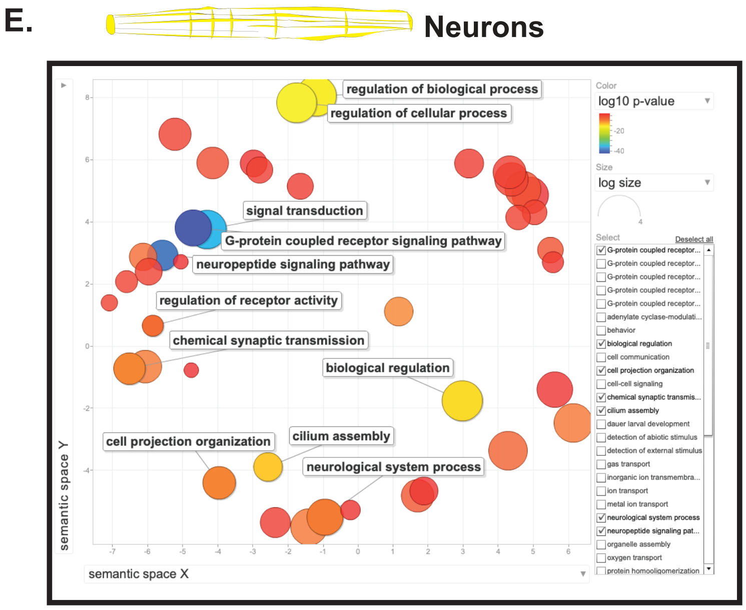


**Figure S6: GO analysis visualized by Revigo of larval tissue specific microarray data from Spencer, et al.** Sematic graphs of GO analysis generated by GOrilla (Eden *et al.* 2009) and visualized by Revigo (Supek *et al.* 2011) of microarray from larval Muscle (BWM, body wall muscle) (**A**) Intestine (Int) (**B**), Hypodermis (Hyp) (**C**), Excretory cells (Exc) (**D**), or Neurons (**E**) (Spencer *et al.* 2011).
