## Supplementary material for "WormCat: an online tool for annotation and visualization of *Caenorhabditis elegans* genome-scale data": Figure S7

Holdorf et al. Figure S7

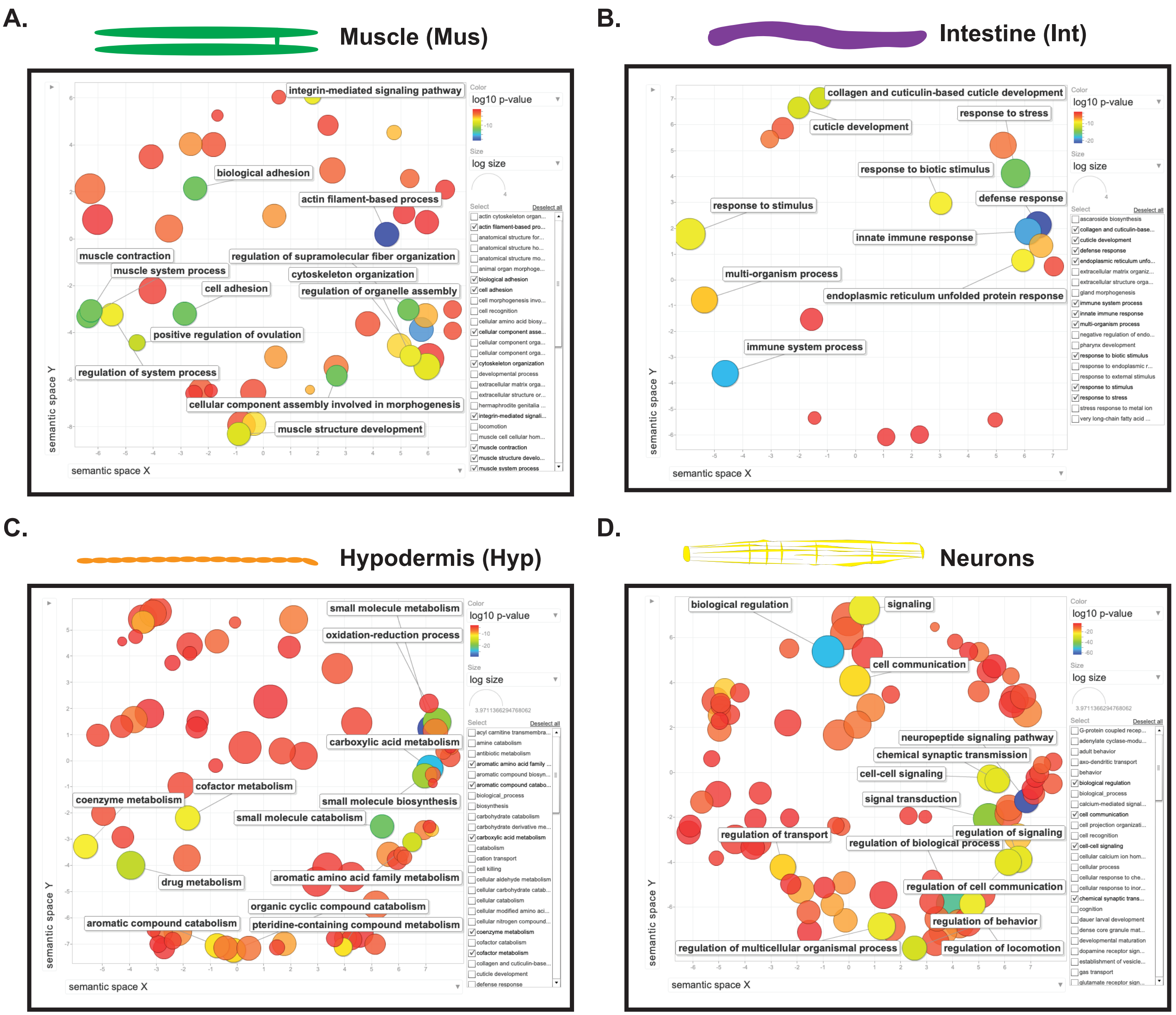


**Figure S7: GO analysis visualized by Revigo of adult tissue specific RNA seq data from Kaletsky, et al.** Sematic graphs of GO analysis generated by GOrilla (Eden *et al.* 2009) and visualized by Revigo (Supek *et al.* 2011) of RNA seq data from adult Muscle (Mus) (**A**) Intestine (Int) (**B**), Hypodermis (Hyp) (**C**), Neurons (**D**) (Kaletsky *et al.* 2018).
