## Supplementary material for "WormCat: an online tool for annotation and visualization of *Caenorhabditis elegans* genome-scale data": Figure S8

Holdorf et al. Figure S8

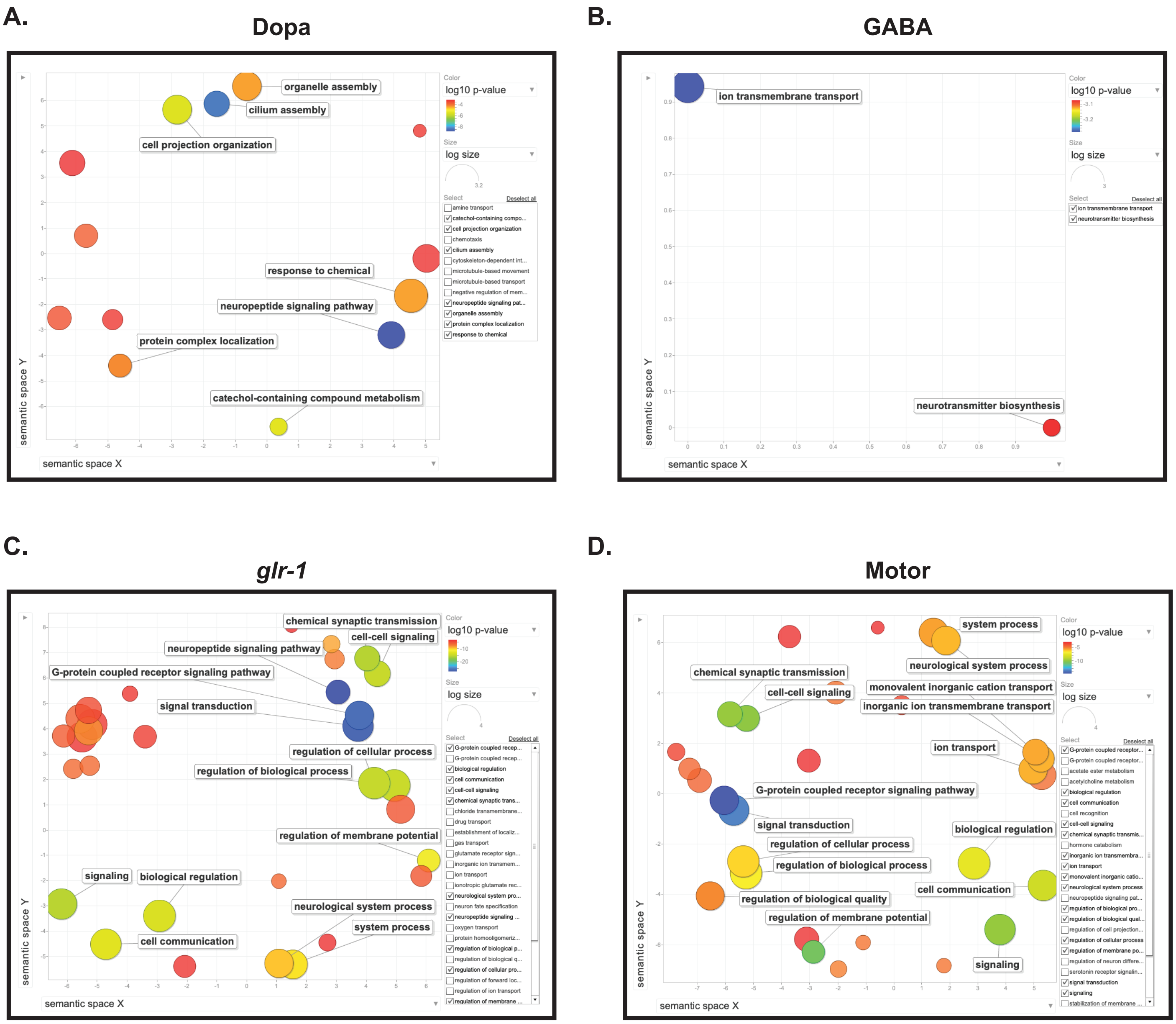


**Figure S8: GO analysis visualized by Revigo of larval neuronal subtype microarray data from Spencer, et al.** Sematic graphs of GO analysis generated by GOrilla (Eden *et al.* 2009) and visualized by Revigo (Supek *et al.* 2011) of microarray from larval dopamenergic (Dopa) (**A**) Gabaergic (GABA) (**B**), *glr-1* expressing (*glr-1*) (**C**), or Class A motor neurons (Motor) (**D**) (Spencer *et al.* 2011).
