## Supplementary material for "WormCat: an online tool for annotation and visualization of *Caenorhabditis elegans* genome-scale data": Figure S9

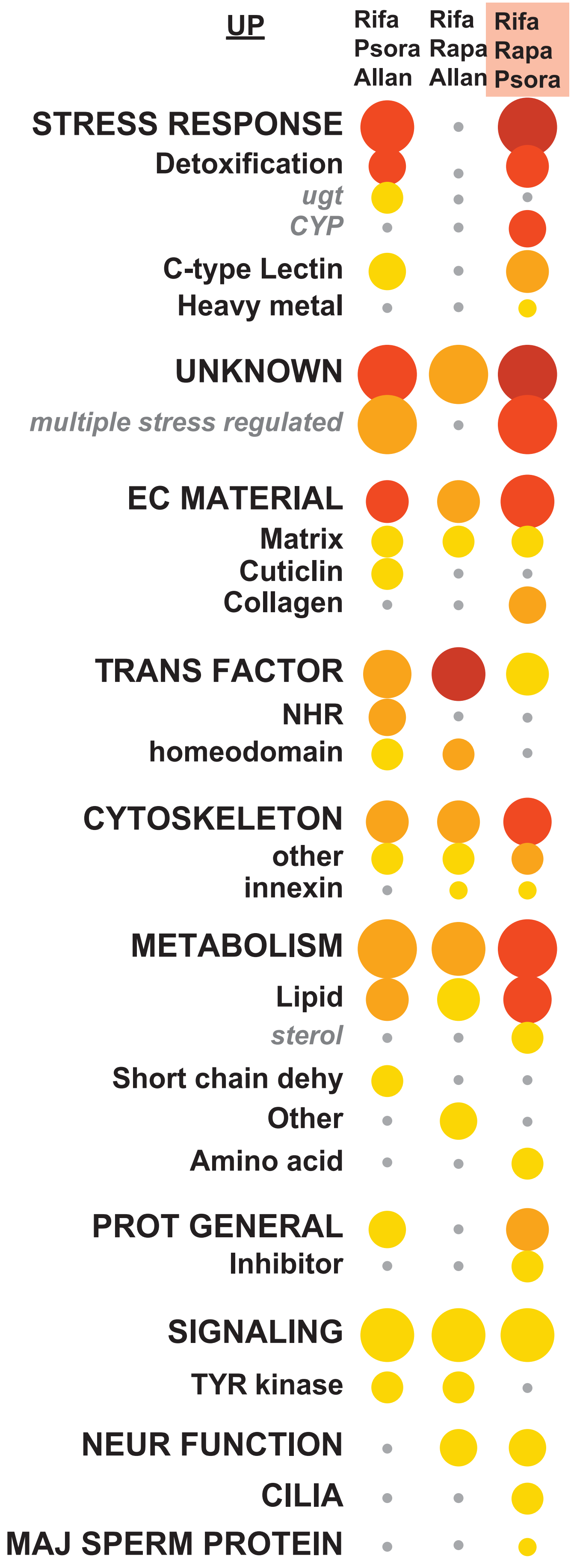


**Figure S9: WormCat analysis of upregulated genes in *C. elegans* treated with triple combinations of lifespan-changing drugs**. Category 1, 2, and 3 analysis of upregulated genes found by RNA-seq from triple-drug combinations (Admasu *et al.* 2018). Pink box denotes drug combination that causes premature death. Allan, allantoin; CYP, Cytochrome P450; EC Material, Extracellular Material; Maj Sperm Protein, Major Sperm Protein; Neur Function; Neuronal Function; NHR, Nuclear Hormone Receptor; Prot General, Proteolysis General; Psora, Psora-4; Rapa, Rapamycin; Rifa, Rifampicin; Short Chain Dehydr., Short Chain Dehydrogenase; Trans Factor, Transcription Factor; TYR kinase, Tyrosine Kinase; ugt, UDP-glycosyltransferase.
