## Supplementary material for "WormCat: an online tool for annotation and visualization of *Caenorhabditis elegans* genome-scale data": Figure S10

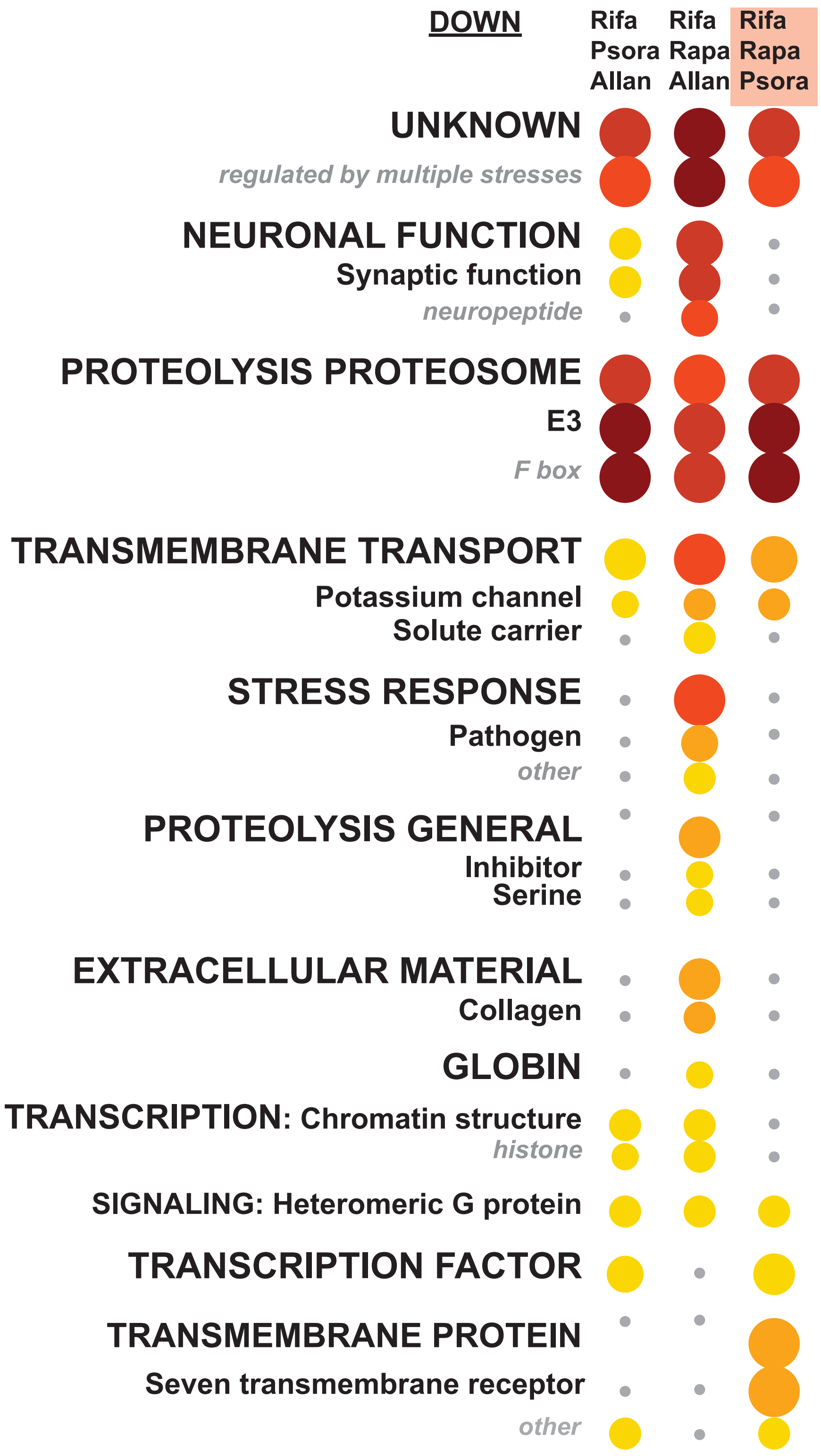


**Figure S10: WormCat analysis of downregulated genes in *C. elegans* treated with triple combinations of lifespan-changing drugs**. Category 1, 2, and 3 analysis of downregulated genes found by RNA-seq from triple-drug combinations (Admasu *et al.* 2018). Pink box denotes drug combination that causes premature death. Allan, Allantoin; Psora, Psora-4; Rapa, Rapamycin; Rifa, Rifampicin.
