## Supplementary material for "WormCat: an online tool for annotation and visualization of *Caenorhabditis elegans* genome-scale data": Figure S11

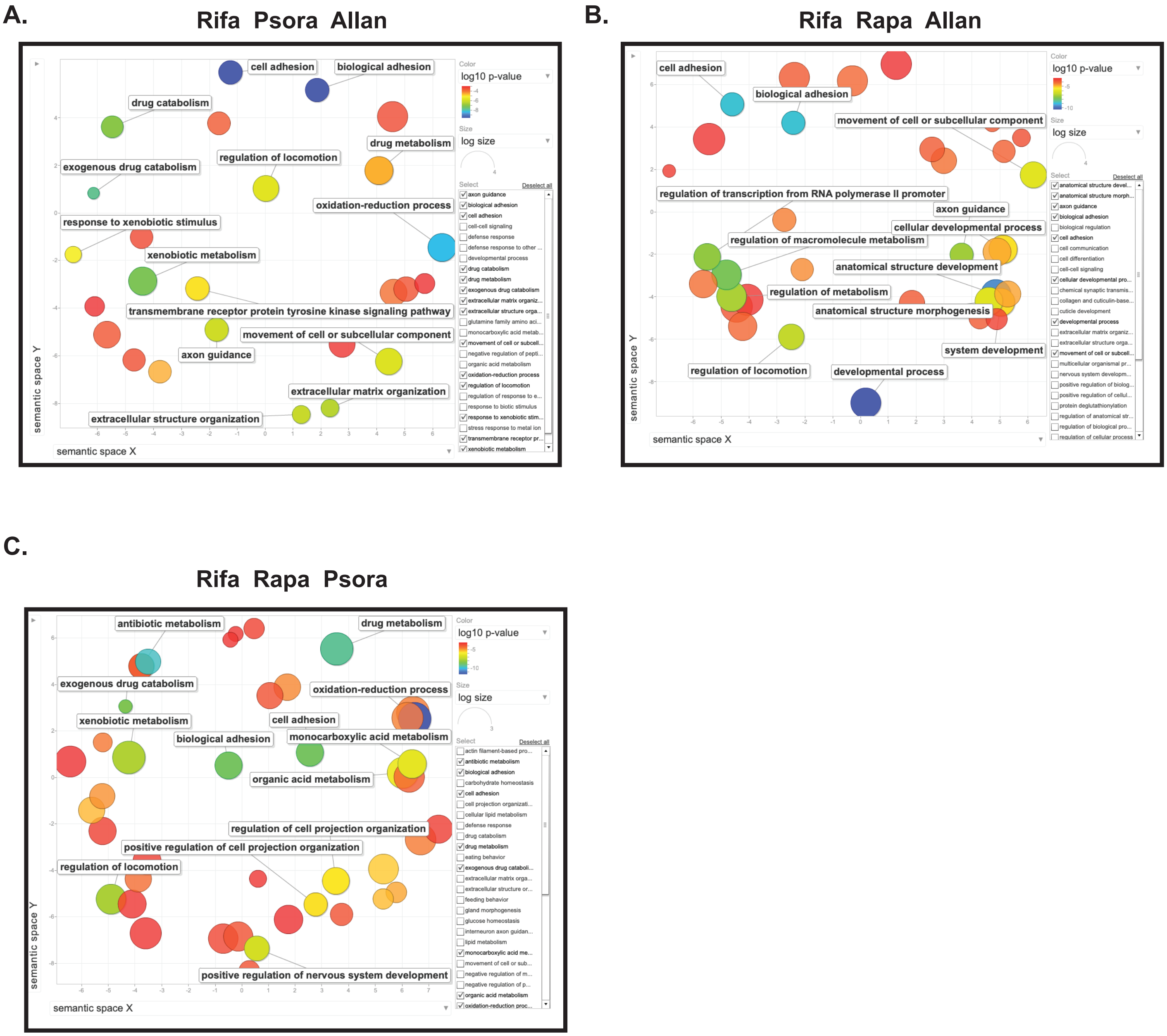


**Figure S11: GO analysis visualized by Revigo of RNA seq data from *C. elegans* treated with triple combinations of lifespan extending drugs from Admasu, et al.** Sematic graphs of GO analysis generated by GOrilla (Eden *et al.* 2009) and visualized by Revigo (Supek *et al.* 2011) of RNA seq data from Rifa, Psora and Allan treated (**A**) Ripa, Rapa, Allan treated (**B**), or Ripa, Rapa, Psora treated (**C**). (Admasu *et al.* 2018).
