## Supplementary material for "WormCat: an online tool for annotation and visualization of *Caenorhabditis elegans* genome-scale data": Figure S12

**Glycogen Low Normal Localization**

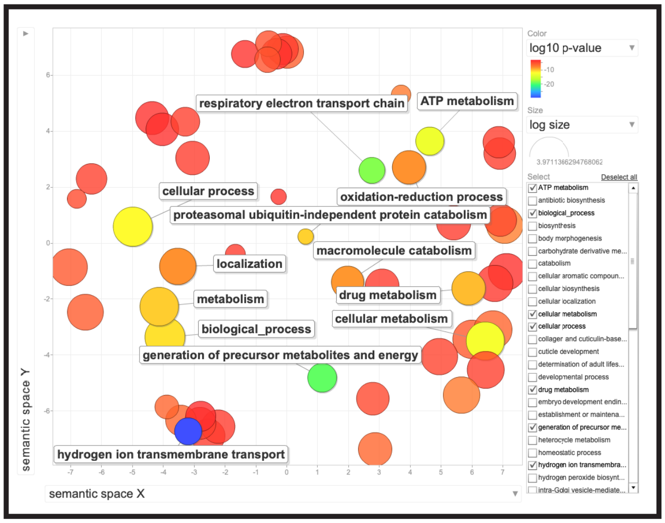


**Figure S12: GO analysis visualized by Revigo of RNA seq data from a *C. elegans* RNAi screen for glycogen storage from LaMacchia, et al.** Sematic graphs of GO analysis generated by GOrilla (Eden *et al.* 2009) and visualized by Revigo (Supek *et al.* 2011) of *C. elegans* showing low glycogen storage in an RNAi screen.
